# Benchmarking Perturbation Tools for the Noncoding Genome

**DOI:** 10.1101/2025.10.21.683665

**Authors:** Han Zhang, Shijie Luo, Xiaofeng Wang, Liquan Lin, Ruipu Liang, Chunge Zhong, Yunhan Zhang, Wenchang Zhao, Zhisong Chen, Xiaoya Liu, Feng Chen, Ning Sun, Jialiang Huang, Teng Fei

## Abstract

Deciphering the functionality of the noncoding genome which includes important *cis*-regulatory elements (CREs) and transcribed noncoding RNA genes remains technically challenging. Here, using massively parallel genetic screening, we systematically benchmark the performance of five representative loss-of-function perturbation tools, including single guide RNA (gRNA) mediated SpCas9 cleavage or CRISPR interference, and paired gRNA (pgRNA) involved dual-SpCas9, Big Papi (paired SpCas9 and SaCas9) or dual-enAsCas12a fragment deletion methods, in decoding the roles of the noncoding genome. For targeting CREs such as enhancer, dual-SpCas9 outperforms other methods with superior efficiency of destroying functional genomic regions. For perturbing noncoding RNA genes, in addition to dual-SpCas9, other RNA-targeting methods such as RNA interference are recommended to discriminate transcript-dependent or -independent roles. A deep learning model DeepDC with associated web server is built to facilitate optimal dual-SpCas9 pgRNA design for efficiently deleting a genomic fragment. Together, our work provides practical guidance on selecting appropriate loss-of-function tools to resolve the functional complexity of the noncoding genome.

## Introduction

Only less than 2% of the human genome encodes proteins and the vast of genomic regions are noncoding^1, 2^. Many important *cis*-regulatory elements (CREs) (e.g., promoter, enhancer, insulator or transcription factor (TF) binding site) and functionally transcribed sequences (e.g., short or long noncoding RNAs (lncRNAs)) are embedded in the noncoding genome^3^. These genetic units not only shape the structural architecture of the genome, but also dictate intricate regulatory and/or effector functions along with protein-coding genes. Many computational efforts have been devoted to annotating or prioritizing potentially functional noncoding regions or genetic variants^4, 5^, however, it remains challenging to experimentally delineate the exact roles of specific noncoding units. Compared to targeting protein-coding genes, the difficulty of perturbing noncoding elements in a loss-of-function manner mainly lies in that (1) the precise demarcation and annotation of a functional noncoding unit at nucleotide resolution is still a tough task; (2) the perturbation tools to precisely disrupt a noncoding unit in the genome are relatively fewer and less efficient; and (3) the noncoding elements are better tolerant to short insertions and deletions (indels) or genetic/epigenetic alterations at a small genomic scale (e.g., several or tens of nucleotides), which precludes us from observing appreciable outcomes or phenotypes even upon a successful disruption at genomic level.

The advent of clustered regularly interspaced short palindromic repeats (CRISPR) based genome editing technology provides efficient perturbation tools for interrogating the noncoding genome^6^. For example, the programmable Cas9 or Cas12 nucleases can generate indels at target genomic loci at a high frequency when associated with a corresponding target-matched guide RNA (gRNA)^6, 7^, thereby disrupting the core DNA sequences at a small scale and possibly the related functions of the noncoding elements^8–15^. Compared to introducing one cut site by a single gRNA (sgRNA), the use of a paired gRNA (pgRNA) strategy with corresponding nucleases enables researchers to create a dual-cut to delete a relatively larger fragment at the target genomic loci^10, 16–23^, which is more likely to produce stronger phenotypes or functional consequences upon perturbation. Moreover, the epigenome editing tools such as CRISPR interference (CRISPRi) or CRISPR activation (CRISPRa) using nuclease-dead Cas9 (dCas9) fusion with transcriptional repressors (e.g., KRAB or KRAB-MeCP2) or activators (e.g., VP64 or p300) or more diverse chromatin modifiers allow varied perturbation patterns and directions to interrogate the functionality of target noncoding loci^24–33^. Nonetheless, loss-of-function still represents a major perturbation way to define the roles of a target genetic element. Although multiple loss-of-function tools exist for perturbing the noncoding genome, it remains puzzling how well these tools perform compared to each other and how to select optimal tools to target specific noncoding elements in the genome.

Here, using high-throughput CRE and lncRNA screens, we systematically benchmark the performance of five typical loss-of-function tools, including sgRNA-mediated SpCas9 knockout (KO) or CRISPRi system, and three pgRNA-involved dual-SpCas9, Big Papi (paired SpCas9 and SaCas9) or dual-enAsCas12a fragment deletion platforms, in functionally perturbing the noncoding genome. Importantly, a deep learning model and associated web server are developed to guide the optimal pgRNA design of dual-SpCas9 system for efficiently deleting a genomic fragment of interest.

## Results

### Building loss-of-function platforms to perturb the noncoding genome

We chose five popular loss-of-function perturbation tools that target the noncoding genome for the benchmark analysis, including (1) conventional SpCas9-based CRISPR KO with a sgRNA; (2) CRISPRi with a sgRNA; (3) dual-SpCas9 double-cut with a pgRNA (a pair of two gRNAs); (4) Big Papi (or Papi for short) double-cut with a pgRNA; and (5) dual-enAsCas12a double-cut with a pgRNA^10, 34–36^. We firstly constructed lentiviral expression plasmids for the five systems (see Methods) (Fig. 1a). The expression levels of different effectors (Cas proteins or their fusion derivatives) across multiple systems were gauged in target cells by controlling the transduced lentiviruses at a low multiplicity of infection (MOI) (< 0.3) before puromycin selection (Extended Data Fig. 1a), thereby maintaining equally low copies of each transduced effector. Using a pre-tested case of known functional noncoding regions for *CUL3* gene regulation in human melanoma A375 cells^15^, we confirmed that all the five systems worked well in decreasing *CUL3* expression when using either sgRNAs or pgRNAs to perturb targeted CTCF binding site, intron or 5’-UTR regions (Fig. 1b). Interestingly, pgRNA-mediated fragment deletion methods caused significantly higher effect size than sgRNA-dependent approaches (Fig. 1c). As pgRNA-involved fragment deletion requires simultaneous cut by two sgRNAs, the dual-cut efficiency is expected to be lower than single-cut and is critical for successful perturbation. Next, we explored the dual-cut efficiencies of the three pgRNA methods on genomic DNA for multiple target loci that were randomly chosen with high or low chromatin accessibility based on ATAC-seq signal in human lung cancer A549 cells. Multiple pgRNAs were designed to delete either long or short fragment for dual-SpCas9, Papi and dual-enAsCas12a systems. The dual-cut efficiency for a given pgRNA of corresponding perturbation system was quantified using the band intensities of PCR-amplified samples after electrophoresis (Fig. 1d and Extended Data Fig. 1b-f). Among the tested 36 cases, we did observe apparent but differential levels of fragment deletion from different pgRNAs by each perturbation system (Fig. 1d and Extended Data Fig. 1b-f), indicating that all the tools were working but the performance of pgRNAs varied a lot. In addition, we found a generally higher editing efficiency for short fragment (∼200 bp) deletion than that of long target (∼500 bp) (Fig. 1e). The target sites located in open chromatin regions tended to be edited more efficiently than those in closed chromatin states (Fig. 1f). Furthermore, dual-SpCas9 seemed slightly superior to the other two systems especially for short fragment deletion in open chromatin using the limited test sites (Fig. 1e,f). Altogether, these results clearly proved the validity of these perturbation platforms in targeting the noncoding genome, but their performance varied for different targets and choices of gRNAs.

**Fig. 1.**
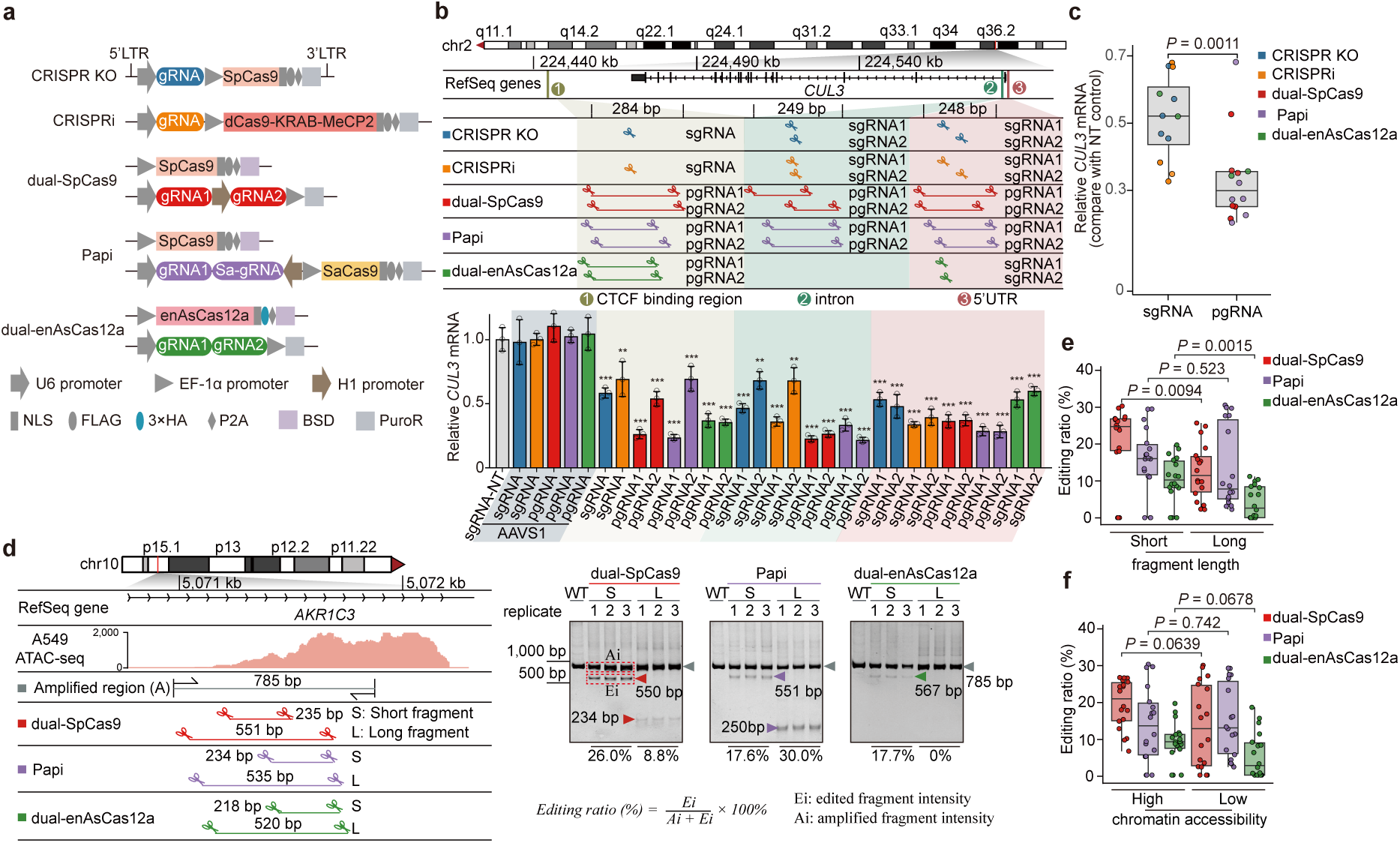
Five typical loss-of-function tools to perturb the noncoding genome. **a**, Schematic of constructed vector architecture for the five indicated perturbation platforms. **b**, Targeted perturbation of the three selected CREs with 5 perturbation platforms in A375 cells for their regulatory functions on *CUL3* gene expression as determined by RT-qPCR. Compared to corresponding *AAVS1*-targeting control, one-way ANOVA. \**P* < 0.05, \*\**P* < 0.01, \*\*\**P* < 0.001. **c**, Comparison of perturbed *CUL3* gene expression change between sgRNA-based CRISPR KO and CRISPRi methods (n = 12 cases) versus pgRNA-mediated dual-cut approaches (n = 14 cases) according to (**b)**. Unpaired two-sample t-test. **d**, Representative dual-cut fragment deletion assay in A549 cells to determine the dual-cut editing efficiency at indicated genomic locus. The designed positions of pgRNAs are shown. Edited samples by corresponding perturbation systems are PCR-amplified across the same region and analyzed by native 10% PAGE in TBE (n = 3 biological replicates). Arrows indicate edited or amplified bands. The calculated editing efficiency is indicated below the gel image. **e**,**f,** Comparison of the dual-cut fragment deletion efficiency of 36 cases (each with 3 biological replicates) by the three perturbation systems shown in (**b**) and (Extended Data Fig. 1b-f) according to either fragment length (short vs. long) (**e**) or chromatin accessibility (high vs. low) (**f**). Two-tailed paired t-tests (n = 18 for each group).

### CRE screening to benchmark multiple perturbation tools

To fully compare the performance of different perturbation systems in targeting CREs such as enhancers, we need to set up a benchmark platform on which all the tools are tested using the same target pool of multiple enhancers with defined functions. However, due to technical difficulty, the number of characterized functional enhancers is still limited so far, especially in a single assay or readout. We managed to find tens of such functional CREs or enhancers that regulate six genes in the 6-thioguanine (6TG)–induced DNA mismatch repair (MMR) pathway and were identified by the Shen group using CRISPRpath strategy in WTC11 iPS cells^37^. Down-regulation of the six 6TG metabolism or MMR genes (*HPRT1*, *MSH2*, *MSH6*, *MLH1*, *PMS2*, and *PCNA*) would counteract the cell death effect elicited by DNA mismatch damage upon 6TG treatment (Extended Data Fig. 2a). The MMR pathway is highly conserved in almost all organisms spanning from bacteria to humans^38^. We further confirmed that knockout of those six genes by CRISPR-Cas9 indeed confer survival advantages over control cells upon 6TG treatment in both A549 and human leukemia K562 cells (Extended Data Fig. 2b-e). Moreover, the identified functional CREs linking to those six genes display similar chromatin openness and acetylated histone H3 lysine 27 (H3K27ac) marks across different cell lines (Extended Data Fig. 2f), further supporting the conservation of these CREs.

We then designed four independent gRNA libraries (one sgRNA library for both CRISPR KO and CRISPRi; three pgRNA libraries for dual-SpCas9, Papi and dual-enAsCas12a) with a total of 13,167 gRNA-effector combinations targeting 69 pre-identified functional CREs for the benchmark purpose (see Methods) (Supplementary Table 1). These independent gRNA libraries were lentivirally transduced into Cas effector-expressing A549 or K562 cells for cell fitness-based CRE or enhancer screens upon vehicle (DMSO) or 6TG treatment (Fig. 2a). The strategies for sgRNA or pgRNA library cloning and next-generation sequencing were detailed in Extended Data Fig. 3a-f (also see Methods). For pgRNA screens, the sequencing reads with shuffled pgRNA sequences due to recombination were removed to obtain the pgRNA counts. The performance of each sgRNA or pgRNA during the screening was quantified by a sigmoid normalization score derived from the log2 fold-change (LFC) values for gRNA counts between the endpoint and Day 0 of the screens (see Methods; Supplementary Table 2). Good correlation was observed between replicates (Extended Data Fig. 4a,b), indicating a high reproducibility of our screening data. Compared to non-targeting or non-functional control gRNAs, a profound portion of sgRNAs or pgRNAs targeting the proximal (< 1 kb to Transcription Start Site (TSS)) or mid-range (between 1-25 kb) enhancers but not distal (> 25 kb) enhancers were positively selected with high sigmoid score upon 6TG treatment (Fig. 2b), suggesting the validity of the screening systems. Moreover, using multiple individual sgRNAs or pgRNAs in different perturbation systems, we further proved that the sigmoid score indeed correlated well with cell fitness phenotype or target gene expression in A549 cells (Fig. 2c,d). Altogether, these results corroborated a high quality of our CRE screens which would be appropriate to benchmark different perturbation tools.

**Fig. 2.**
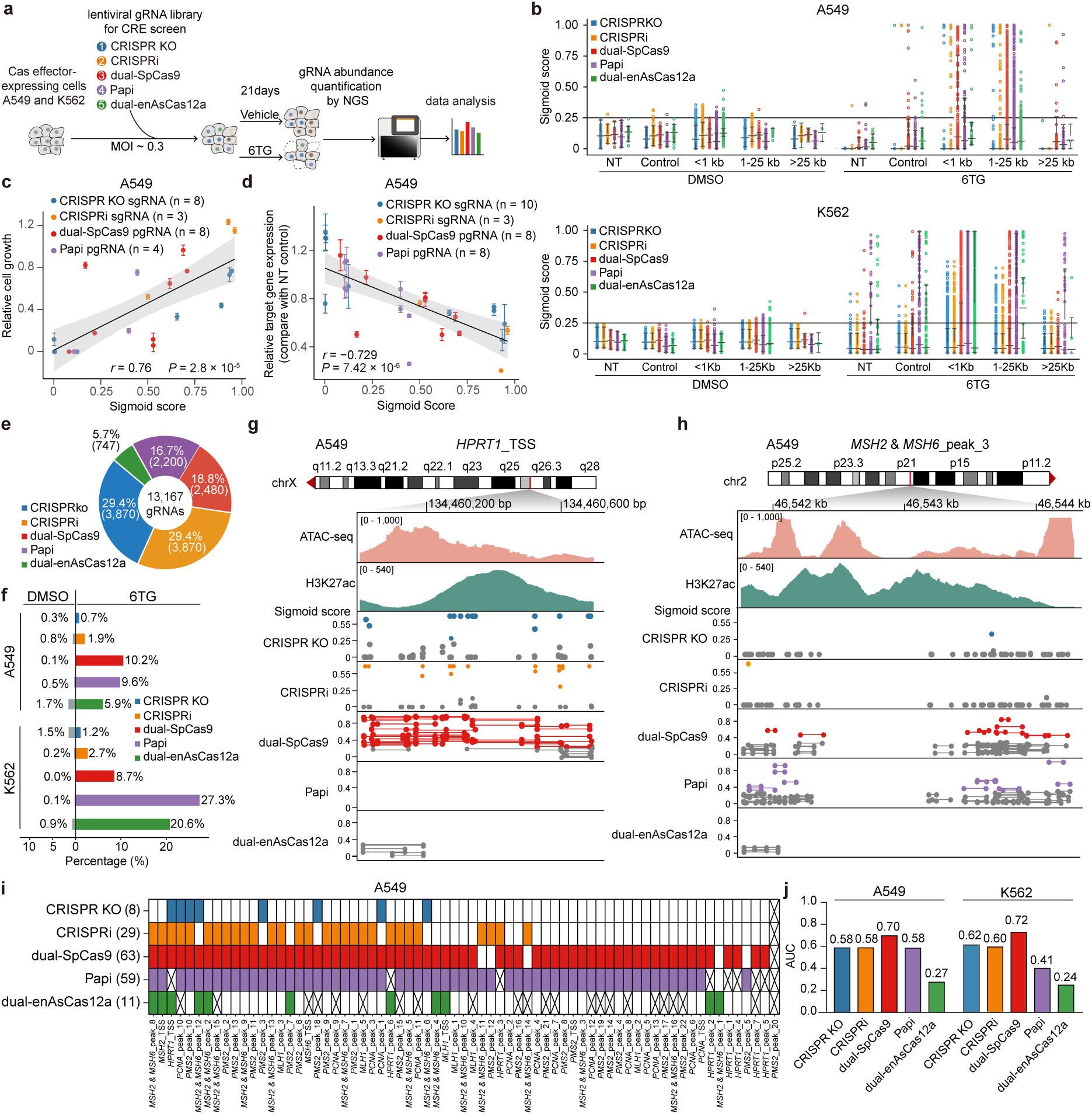
CRE screens to benchmark multiple perturbation tools. **a**, Schematic of CRE or enhancer screening process. **b**, Normalized sigmoid score distribution of sgRNAs or pgRNAs in the A549 and K562 cell screens according to the distance of targeted loci from TSS in DMSO or 6TG treatment conditions. NT: non-targeting gRNAs, Control: gRNAs targeting *AAVS1*, *CCR5* or *ROSA26* sites. **c**, Scatter plot showing the relationship between sigmoid score and phenotypic change (relative cell growth, compared to *AAVS1* control, mean ± SEM of 3 biological replicates) after individual sgRNA or pgRNA perturbation in A549 cells. Each data point represents one gRNA for indicated perturbation platforms. Pearson’s *r* and associated *P* value are shown. **d**, Scatter plot showing the relationship between sigmoid score and relative target gene expression for each indicated gRNA and perturbation systems in A549 cells (compared to NT control pgRNA, mean ± SEM, n = 3). Pearson’s *r* and two-tailed associated *P* value are indicated. **e**, The gRNA number and percentage of 5 platforms in the benchmarking CRE library. **f**, The percentages of positively selected sgRNAs or pgRNAs of different perturbation platforms in 6TG or DMSO condition during A549 and K562 cell CRE screens. **g,h**, Representative screening results showing the detailed gRNA performance (sigmoid score in 6TG) in the CRE screen for the indicated *HPRT1_*TSS (**g**) or an enhancer (**h**) loci in A549 cells. **i**, Heatmap showing the functional enhancers identified in A549 cells by each perturbation platform. The crossed grid indicates that no valid gRNAs were designed for the corresponding CRE. **j**, The area under the ROC curve (AUC) comparison for gRNA performance of all five perturbation platforms in A549 or K562 cells.

The number of designed sgRNAs or pgRNAs for different SpCas9-based systems were similar (Fig. 2e). However, the designable pgRNAs for dual-enAsCas12a were much fewer (Fig. 2e), due to more stringent restriction of its favorable protospacer adjacent motif (PAM) (TTTV)^36^. The percentages of positively selected sgRNAs or pgRNAs (sigmoid score > 0.25) upon 6TG treatment dramatically increased compared to those in vehicle condition (Fig. 2f). Interestingly, the three pgRNA-involved perturbation systems generally displayed higher percentages of positively enriched gRNAs than sgRNA-based CRISPR KO or CRISPRi system (Fig. 2f). These results indicated that dual-cut fragment deletion methods are more likely to produce functional consequences than single site-targeting approaches for CRE perturbation. From the gRNA profiles of several representative CREs, we observed differential performances of these perturbation tools and a large portion of gRNAs were usually not working (Fig. 2g,h and Extended Data Fig. 4c,d), further substantiating the difficulty of perturbing the noncoding genome. Therefore, functional CREs in the screens were defined for each perturbation system according to the following arbitrary criteria: (1) a qualified gRNA with sigmoid score^6TG^ > 0.25 & (score^6TG^ - score^DMSO^) > 0.1; and (2) at least 2 qualified gRNAs exist for the target CRE. Under the same criteria, differential number of functional CREs were identified by each perturbation platform, with dual-SpCas9 system capturing the most functional enhancers in both A549 and K562 cells (Fig. 2i and Extended Data Fig. 4e; Supplementary Table 2). To compare the accuracy of gRNAs of each perturbation system, we used high-confidence functional CREs (identified in at least 3 perturbation systems) and non-functional controls as references, and curated the gRNA performance by a Receiver Operating Characteristic (ROC) curve analysis. As shown in Fig. 2j, dual-SpCas9 performed the best over other methods in both A549 and K562 cell lines, with the area under the ROC curve (AUC) of 0.70 and 0.72, respectively. These data suggest that dual-SpCas9 is the preferred tool, among others, to functionally perturb the CREs or enhancers.

### Interrogating noncoding RNA genes using different perturbation systems

Disruption of promoter regions by CRISPR knockout or blocking transcription via CRISPRi represent typical strategies to functionally perturb the ncRNA genes at the genomic level^19, 39, 40^. To explore the capabilities of different perturbation systems in targeting the ncRNAs, we constructed one sgRNA library (for both CRISPR KO and CRISPRi) and three pgRNA libraries (for dual-SpCas9, Papi and dual-enAsCas12a) with a total of 17,257 gRNA-effector combinations targeting ∼200 lncRNAs that were highly expressed in human lung carcinomas (Extended Data Fig. 5a; Supplementary Table 3; see Methods). Multiple independent screens were performed in human lung cancer PC-9 cells using cell fitness as the readout to interrogate functional lncRNAs regulating cell growth or survival (Fig. 3a). The gRNAs were quantified by high-throughput sequencing and the recombination reads were removed for pgRNAs before proceeding to downstream analysis. Replicates correlated well (Extended Data Fig. 5b), and the control gRNAs targeting the coding regions of core essential genes were negatively selected except for CRISPRi system whose gRNAs should work around the TSS rather than the coding regions (Fig. 3b). These data indicated a high quality and validity of the screen systems. MAGeCK algorithm was employed to determine the functional lncRNAs by a transformed RRA (tRRA) score^41^. A higher positive score for a given lncRNA indicates stronger positive selection upon perturbation, while a lower negative score means stronger negative selection.

**Fig. 3.**
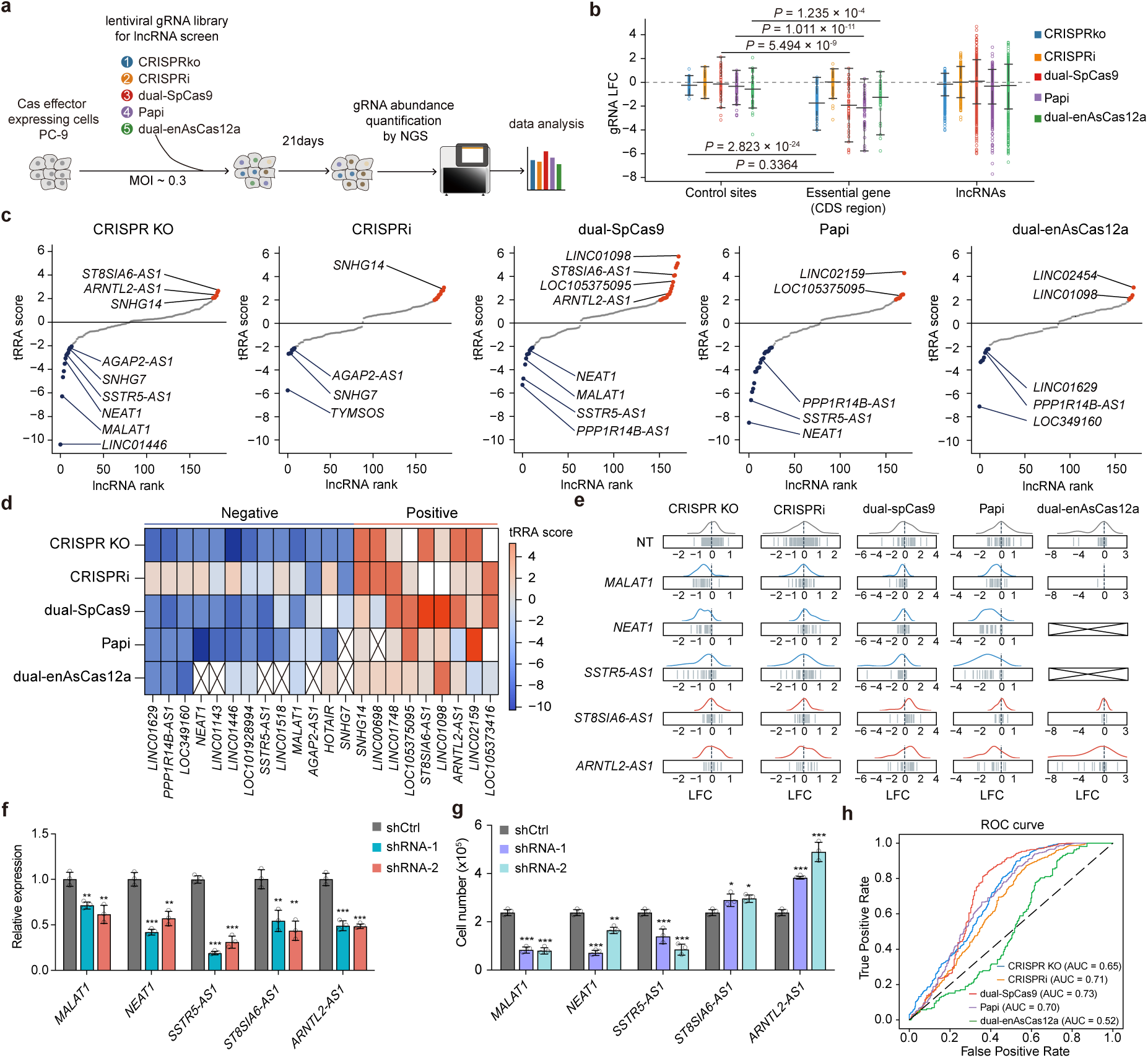
Benchmarking perturbation tools for interrogating ncRNA genes. **a**, Schematic of the lncRNA screening workflow using five perturbation platforms. **b**, Log₂ fold change (LFC) of individual sgRNA or pgRNA counts (endpoint vs. D0) in control, essential genes, and lncRNA groups in PC-9 cells. Control targets including *AAVS1*, *CCR5*, and *ROSA26*; essential gene targets covering CDS regions of *RPS19, RPS20, RPL7, EEF1A1,* and *EEF2*. **c**, Highlights of several functional lncRNAs identified by each perturbation platform in PC-9 cells. **d**, Heatmap showing the functional lncRNA landscape colored by tRRA scores across five perturbation systems in PC-9 cells. The crossed grid indicates that no valid gRNAs were designed for the corresponding lncRNA. **e**, LFC deviation of selected targets during the lncRNA screening in PC-9 cells. **f**, Relative expression levels of selected lncRNAs following shRNA-mediated knockdown in PC-9 cells. Mean ± SD, n = 3, relative to vector control, one-way ANOVA, \**P* < 0.05; \*\**P* < 0.01; \*\*\**P* < 0.001. **g**, Cell growth of PC-9 cells 7 days after shRNA-mediated knockdown of selected lncRNAs. Mean ± SD, n = 3 biological replicates, compared to vector control, one-way ANOVA, \**P* < 0.05; \*\**P* < 0.01; \*\*\**P* < 0.001. **h**, ROC curve analysis comparing the gRNA performance of all five perturbation platforms.

Using a uniform cutoff (tRRA score > 2 or < −2), tens of cell growth-related lncRNAs were identified in PC-9 cells by each perturbation tool (Supplementary Table 4). Several known lncRNAs, such as *NEAT1* and *MALAT1*, were successfully identified in more than two perturbation systems (Fig. 3c), which agree with their reported functions in lung cancer^42, 43^. The selection directions of high-confidence functional lncRNAs (identified in at least 2 perturbation systems) were largely accordant across different systems (Fig. 3d). The gRNA performance for representative lncRNAs or control targets during the screens was shown in Fig. 3e. To validate the functions of these lncRNAs, we employed RNA interference (RNAi), another orthogonal perturbation tool at the RNA level, to individually knockdown these lncRNAs in PC-9 cells (Fig. 3f). Indeed, the cell growth effects upon the lncRNA knockdown with two independent short hairpin RNAs (shRNAs) were concordant with those shown in the screens, with *ST8SIA6-AS1* and *ARNTL2-AS1* knockdown promoting cell fitness while *MALAT1*, *NEAT1*, *SSTR5-AS1* knockdown decreasing cell growth (Fig. 3g), further corroborating the accuracy of our perturbation screens. Using functionally confirmed lncRNAs (supported by the literatures and also identified in our screens) and non-functional controls as references, we performed ROC curve analysis and found that the pgRNAs in dual-SpCas9 system displayed the highest accuracy (AUC = 0.73) over the other tested tools in functionally targeting the ncRNA genes at the genomic level (Fig. 3h). This finding is consistent with that obtained from CRE screens (Fig. 2j).

Notably, almost no single functional lncRNA could be identified by all the five perturbation systems (Fig. 3d; Supplementary Table 4), suggesting that each tool has its intrinsic limitation. As all these five systems are DNA-targeting, there might be tiny chances that (1) the perturbation causes genotoxicity effects especially for the DNA-cutting tools; (2) the targeted TSS region according to the standard RefSeq annotation is not the genuine TSS of targeted ncRNA transcripts or isoforms in specific cell contexts; (3) the targeted DNA region (TSS of the targeted ncRNA gene) might simultaneously have additional regulatory functions (e.g., as an enhancer) on other genes. If any of the above side-effect or off-target scenarios happens, the perturbation on the genomic DNA level would not faithfully reflect the RNA transcript’s function of the targeted ncRNA gene. Indeed, we managed to locate one such case in our lncRNA screens. The lncRNA gene *LINC01098* showed consistently positive selection in multiple DNA-acting perturbation systems (Fig. 3d and Extended Data Fig. 5c), however, RNAi-mediated transcript knockdown exhibited significantly reduced cell fitness effects (Extended Data Fig. 5d,e). Therefore, RNA-acting perturbation tool is necessary to discriminate the RNA-dependent or -independent roles of interrogated ncRNA gene.

### A deep learning model DeepDC for predicting pgRNA-mediated fragment deletion efficiency by dual-SpCas9

Our systematic CRE and lncRNA screens suggest that dual-SpCas9 is superior to other perturbation tools in targeting the noncoding genome. Nevertheless, how to design efficient gRNA pairs to increase the likelihood of successful dual-cut and fragment deletion remains elusive. To solve this problem, we assembled a comprehensive dataset with the noncoding genome perturbation results from 154,457 dual-SpCas9 pgRNAs targeting CREs or ncRNA genes across multiple cell lines by integrating our data in current study with previously published screens (Fig. 4a). After rigorous data refinement by re-running MAGeCK and direction-specific sigmoid normalization, the top and bottom 10% of pgRNAs were selected and partitioned into training and test sets (7:3 split) (Fig. 4b). Building on this resource, we developed DeepDC (a <u>Deep</u> learning model for predicting pgRNA activity on <u>D</u>ual-<u>C</u>ut fragment deletion by dual-SpCas9), which is a predictive framework that combines XGBoost with multi-dimensional features, including target fragment length, GC content, sgRNA efficiency index, epigenetic signals, and fragment-level scores extracted by the sequence-sensitive agent Hyena^44^ (Fig. 4b). As the first deep learning model specifically designed for predicting pgRNA efficiency of dual-cut fragment-level editing, DeepDC showed superior performance compared to several existing single-cut editing prediction tools or the Random Forest algorithm across multiple evaluation metrics (AUC, Accuracy, Precision, Recall, and F1 scores) under 5-fold cross validation (Fig. 4c). Intriguingly, gradient-based analyses revealed that the fragment (the sequences between the dual-cut) related features (particularly fragment length and Hyena score) showed more representation than the sgRNA property (harmonic mean of predicted efficiency scores of the two paired sgRNAs) or epigenetic features of the target region among the high-dimensional determinants of DeepDC model (Fig. 4d). This result suggests that the dual-cut fragment editing efficiency should not be simply deduced from the sgRNA cutting efficiency on either end. The re-ligation efficiency of two chromatin ends by the DNA repair machinery after dual-cut might also be important, which could be severely affected by the fragment length, local chromatin sequence or epigenetic states.

**Fig. 4.**
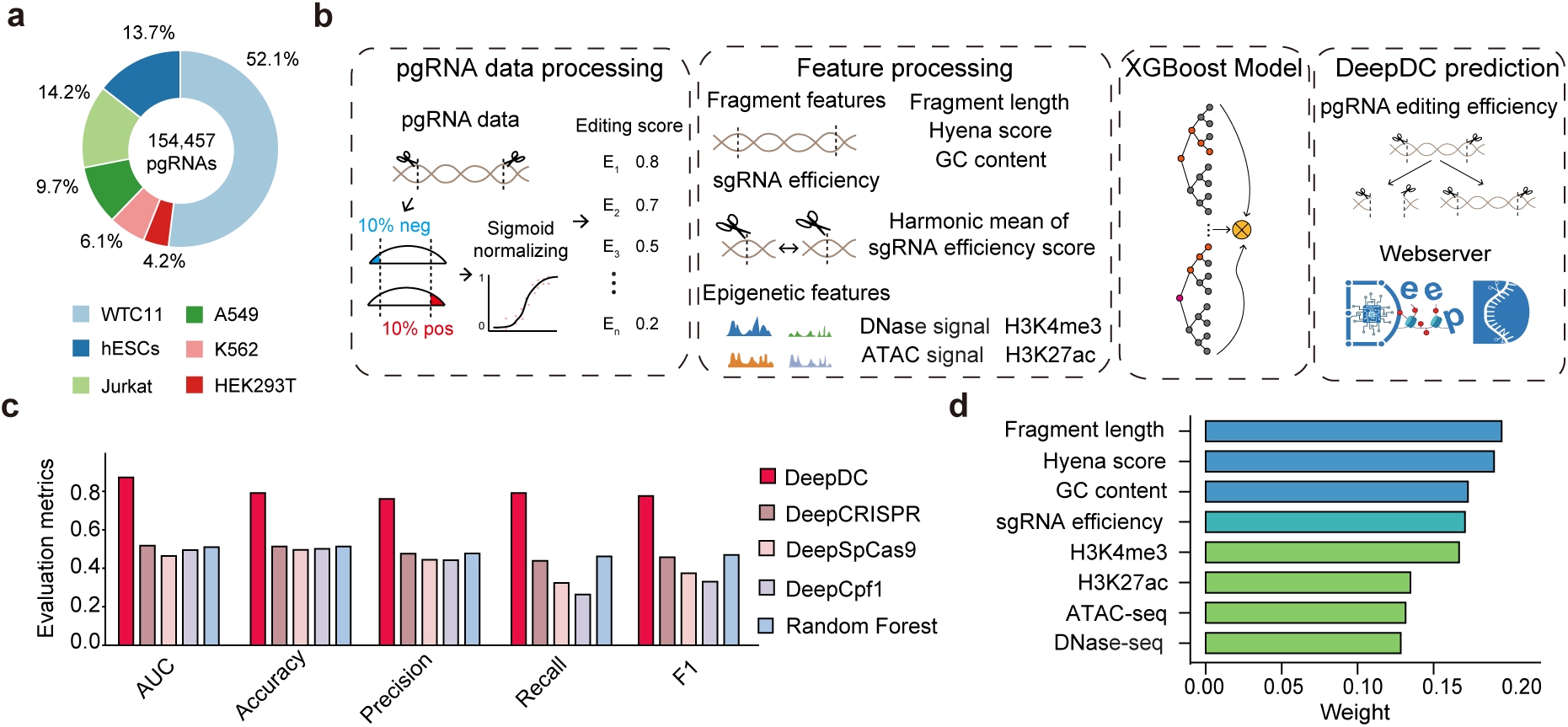
A deep learning model DeepDC for dual-cut fragment deletion by dual-SpCas9. **a**, Data collection summary of dual-SpCas9 pgRNA perturbation events from public studies and our screenings catalogued by the cell types. **b**, A schematic overview of DeepDC model development. **c**, Performance comparison of DeepDC against existing sgRNA efficiency predictors or machine learning methods, including DeepCRISPR, DeepSpCas9, DeepCpf1, and Random Forest, evaluated using five metrics: AUC, Accuracy, Precision, Recall, and F1-score. **d**, Average feature weights assigned by the DeepDC model. Fragment related features (length, GC content, Hyena score), sgRNA efficiency, and epigenetic features (H3K4me3, H3K27ac, ATAC, and DNase I signals) collectively contribute to the prediction of dual-SpCas9 pgRNA efficacy.

To facilitate the user’s application, we built a user-friendly webserver for DeepDC (Extended Data Fig. 6; https://deepdc.huanglabxmu.com/static/index.html) along with a standalone version (https://github.com/xmuhuanglab/DeepDC) to facilitate the design of optimal dual-SpCas9 pgRNAs. The input of DeepDC was a genomic fragment of interest for perturbation, the output was a set of candidate pgRNAs with predicted DeepDC score ranging from 0 to 1, with a higher score indicating a better chance of successful dual-cut fragment editing.

### Applying DeepDC model for enhancer characterization

To showcase the robustness and application of the DeepDC model, we chose a representative case of enhancer characterization, in which the expression of β-like globin genes (e.g., *HBE1*, *HBG1/2* and *HBB*) could be repressed when disrupting a cluster of specific enhancers including DNase I hypersensitive sites HS1 and HS2^28^. Using the DeepDC model, we designed a bunch of pgRNAs with high (> 0.5) or low (< 0.2) predicted scores against HS1 and HS2 (Fig. 5a). Next, we individually transduced these pgRNAs into SpCas9-expressing K562 cells by lentiviruses, and determined the dual-cut efficiency of each pgRNA pair (Fig. 5b). As expected, the pgRNAs with high DeepDC scores displayed significantly higher editing (dual-cut) efficiency than those with low scores (Fig. 5c). Consistently, high-score pgRNAs caused more significant repression of the β-like globin gene expression than those low-score pgRNAs (Fig. 5d,e). These results further proved the robustness of our DeepDC model for improved pgRNA design and dual-cut fragment deletion in functionally characterizing the noncoding genome.

**Fig. 5.**
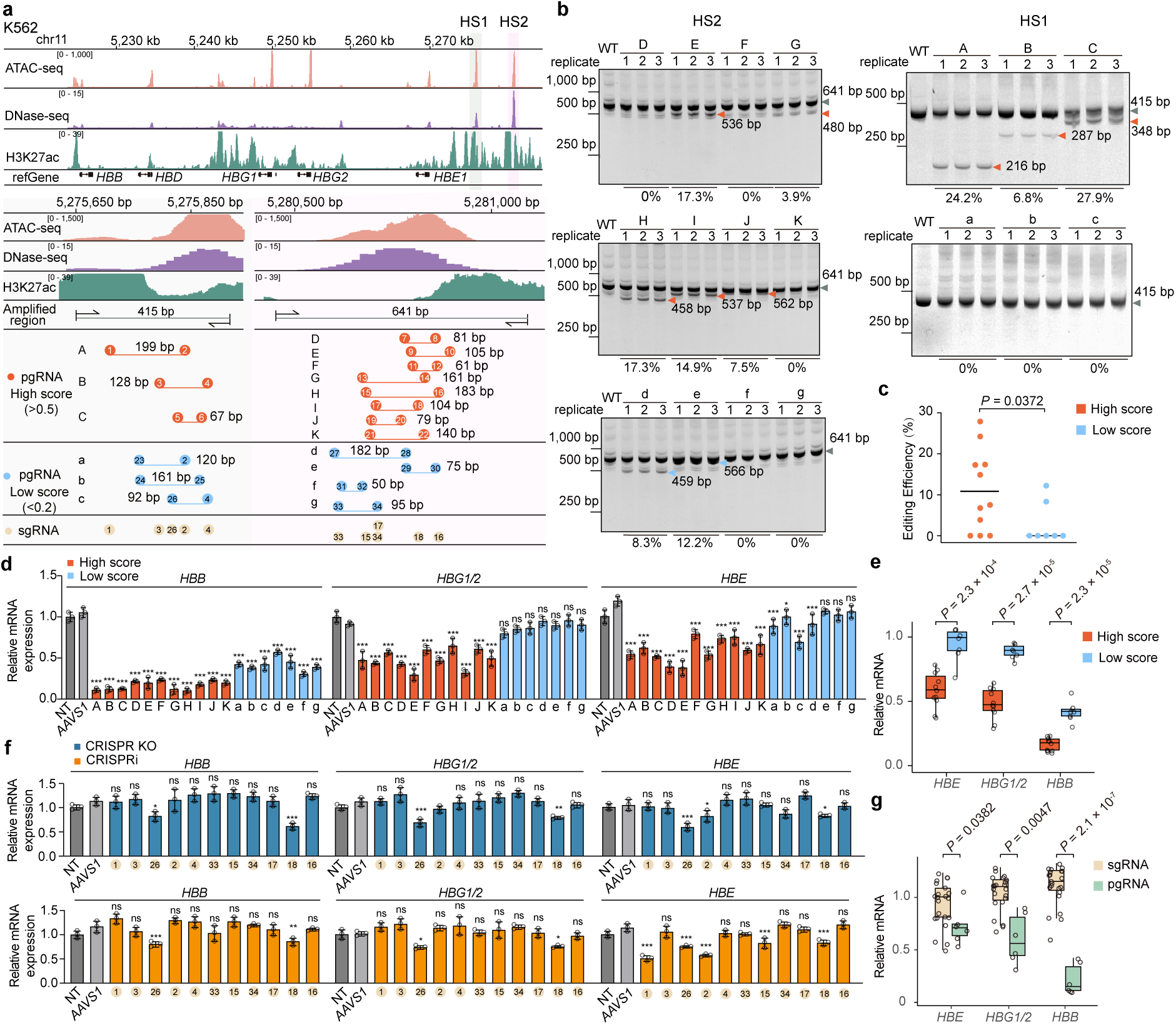
Validation of the DeepDC model in pgRNA selection for enhancer characterization. **a**, Schematic showing the position of DeepDC-scored pgRNAs targeting HS1 and HS2 enhancers which regulate the β-like globin genes (e.g., *HBE1*, *HBG1/2* and *HBB*). For each region, several high-score (n = 11) and low-score (n = 7) pgRNAs are designed. **b**, TBE gel electrophoresis showing the pgRNA editing results for each indicated pgRNA in K562 cells. Arrows indicate edited or amplified bands. **c**, Comparison of pgRNA editing efficiency between high-score and low-score pgRNA groups. Each dot represents an average editing ratio from three biological replicates for one pgRNA evaluation. One-tailed unpaired t-test. **d**, Relative mRNA expression of β-globin genes in K562 cells after perturbation with high- or low-score pgRNAs according to DeepDC, normalized to non-targeting pgRNA control. Mean ± SEM, n = 3, compared to pg*AAVS1* control, one-way ANOVA, \**P* < 0.05, \*\**P* < 0.01, \*\*\**P* < 0.001, ns, not significant. **e**, Comparison of β-globin gene expression after dual-SpCas9 pgRNA perturbation between the high-score and low-score pgRNA groups. One-tailed unpaired t-test. **f**, Relative mRNA expression of β-globin genes following perturbation with the pgRNA-matched individual sgRNAs shown in (**a**) by either CRISPR KO or CRISPRi perturbation systems. Mean ± SEM, n = 3, compared to pg*AAVS1* control, one-way ANOVA, \**P* < 0.05, \*\**P* < 0.01, \*\*\**P* < 0.001, ns, not significant. **g**, Direct comparison of β-globin expression after perturbation by sgRNAs (through either CRISPR KO or CRISPRi) versus their matched pgRNAs (through dual-SpCas9). Two-tailed unpaired t-test.

In addition, we also designed several pgRNA-matched sgRNAs and determined the perturbation effects by CRISPR KO and CRISPRi approaches (Fig. 5a). Although more than half of the sgRNAs were derived from high-score pgRNA pairs, only two of them (sgRNA-26 and sgRNA-18) exhibited significant gene repression for all the three tested genes (Fig. 5f). Interestingly, the perturbation effects of sgRNA-26 and sgRNA-18 were consistent between both CRISPR KO and CRISPRi systems (Fig. 5f), indicating the functional conservation of the two sgRNAs. Notably, sgRNA-26 was derived from a low-score pgRNA-c, whereas sgRNA-18 was retrieved from a high-score pgRNA-I (Fig. 5a). This further suggests that the editing efficiency of a pgRNA pair cannot be simply deduced from that of either sgRNA, which strengthens the necessity of DeepDC scoring model for pgRNA selection. Moreover, the phenotypic effects (gene repression) of pgRNA-mediated perturbation were significantly larger than those of sgRNA-dependent approaches (Fig. 5g), further justifying the priority of using dual-SpCas9 to interrogate functional CREs.

## Discussion

Efficient perturbation of a noncoding genomic locus in a loss-of-function manner is necessary to decipher its functionality and causality for complex phenotypes in genetic studies. Here we comprehensively compare five popular CRISPR-based perturbation tools for the loss-of-function usage against CREs and ncRNA genes. Our study not only uncovers varied performance of multiple tools in targeting different types of noncoding units, but also provides practical guidance on appropriate toolkit selection and improved experimental design towards better efficiency and more precise conclusions.

In contrast to protein-coding genes which are usually sensitive to frameshift mutations, noncoding genetic elements might be more tolerant to small indels for appreciable phenotypes. Consistently, we did find that large fragment deletion methods depending on pgRNA-mediated dual-cut tend to identify functional noncoding elements at higher efficiency than sgRNA-mediated CRISPR KO or CRISPRi approaches. Furthermore, among the three double-cut fragment deletion tools, dual-SpCas9 performs the best, which is likely due to its superior DNA cleavage efficiency compared to that of SaCas9 and enAsCas12a. The more stringent PAM restriction for SaCas9 (NNGRR)^45^ and enAsCas12a (TTTV)^36, 46^ further limits the choice of designable number of gRNAs. Therefore, it is advisable to deploy dual-SpCas9 for perturbing CREs such as enhancers or transcription factor binding sites. As an orthogonal or complementary approach, CRISPRi is still highly recommended as it works through dCas9-mediated steric hindrance of obligate binding factors and/or effector domain-elicited transcriptional repression^47^, which avoids the uncontrolled side effects and/or genotoxicity resulting from DNA cleavage. However, the perturbation strength of sgRNA-mediated CRISPRi seems to be comparable to single-cut SpCas9 KO but weaker than double-cut dual-SpCas9 system. This phenomenon was also observed in another recent study that focused on enhancer characterization^32^. The perturbation strength of CRISPRi system might be enhanced via careful selection of best-performing gRNAs, simultaneous targeting using multiplexed gRNAs, or applying other highly efficient CRISPRi effectors.

For transcript-producing noncoding genes such as miRNAs, lncRNAs or other ncRNAs, destroying their genomic DNA with cleavage-based perturbation tools can also work well as they do for CREs in terms of perturbation efficiency. Notably, the DNA cleavage strategy (e.g., deleting promoter, splice sites or specific exons of ncRNA gene) might also affect the perturbation outcomes even for the same ncRNA gene^19, 39, 48^. Still, dual-SpCas9 system outperforms other DNA-acting perturbation methods in our benchmark screens for the lncRNA genes. The perturbation effects of CRISPRi seemed to be generally weaker compared with other DNA cleavage based tools for those high-confidence hits on both negative and positive directions (Fig. 3d). It is unlikely that the difference is due to potential genotoxicity caused by other DNA-cutting systems, as most of the negatively selected high-confidence hits can be functionally supported in lung cancer by other published studies (Supplementary References) (e.g., the well-known essential lncRNAs *NEAT1* and *MALAT1*). We surmise that, compared to functional element deletion, the steric hindrance and/or transcriptional inhibition of CRISPRi apparatus cannot fully block the RNA production especially for those highly-expressed genes, thereby leading to less significant perturbation effects in general. Considering the complicated transcriptional heterogeneity and chromatin context, it remains technically challenging to robustly manipulate the expression of any given gene for such epigenome editors. Hence, additional improvement efforts are needed as stated above to reinforce the perturbation robustness of CRISPRi system. On the other hand, researchers should also be cautious about the results obtained from all these DNA-acting perturbation tools when studying transcript-producing ncRNA genes due to potential side-effects and/or off-targets from DNA-targeting. In such scenario, orthogonal RNA-acting perturbation methods (e.g., RNAi or CRISPR-Cas13) are mandatory to distinguish the transcript-dependent or -independent functions for a given ncRNA gene.

Notably, the comparative analysis of each perturbation system was based on a large-scale measurement with their default or routine parameters and procedures, and therefore the conclusion here was largely statistical or in a general sense. Therefore, for a specific site perturbation at an individual basis, other or additional perturbation systems would be considered with improved operating conditions if the first attempt does not give satisfactory results or seeking for the best effect.

Altogether, the above lessons learned from this benchmark study along with a web-implemented deep learning model DeepDC provide practical guidance on how to efficiently perturb a noncoding CRE or a ncRNA gene in a loss-of-function manner. The combination of these tools with additional cutting-edge techniques^49–57^ such as high-throughput genetic screens, single-cell and spatial multi-omics hold great promise to decoding the mystery of complex noncoding genome and genetic variants at unprecedented speed and depth.

## Methods

### Vector construction

All the plasmids expressing the perturbation systems were assembled and delivered via a lentiviral system. For CRISPR KO, the lentiCRISPRv2 (Addgene, #52961) vector was used. For CRISPRi vector (plenti-dCas9m4-KRAB-MECP2-puro), the plasmid was assembled with XbaI (Takara, #1634) and BamHI (Takara, #1605) digested lentiCRISPRv2 vector, the Cas9-encoding sequence with four amino acid mutations (D10A, D839A, H840A, and N863A), and the KRAB-MeCP2 fragment amplified from dCas9-KRAB-MeCP2 vector (Addgene, #110821) via Gibson assembly. For dual-SpCas9, the pLenti-Cas9-Blast vector (Addgene, #52962) was used to generate the Cas9-expressing cell line, and the plentiGuide-Puro vector (Addgene, #52963) served as the pgRNA expression vector. For Big Papi, the pLenti-Cas9-Blast vector was also utilized to establish the SpCas9-expressing cell line. The pPapi plasmid functioned as both the pgRNA expression and the SaCas9 expression vector. To construct the pPapi vector, lentiCRISPRv2 was digested with XbaI and BamHI to remove the Cas9 coding sequence, which was subsequently replaced with the SaCas9 fragment PCR-amplified from pX601 vector (Addgene, #61592). The assembled SaCas9-expressing lentiCRISPRv2 was then digested with BsmBI (New England Biolabs, R0739L) and EcoRI (Takara, #1611) to remove the filler and gRNA scaffold sequence. Next, a reversed H1 promoter was inserted, restoring two Esp3I recognition sites to facilitate the insertion of gRNAs. For dual-enAsCas12a, the pRDA_174 vector (Addgene, #136476) was used to generate the enAsCas12a-expressing cell line, while pRDA_052 (Addgene, #136474) served as the pgRNA expression vector.

### CRISPR library design

For the enhancer library design, a total of 69 previously identified CREs or enhancer regions associated with six key genes in the MMR pathway (*HPRT1*, *MSH2*, *MSH6*, *MLH1*, *PMS2*, and *PCNA*) were selected for perturbation^37^. For the pgRNA-involved perturbation platforms, the individual gRNAs were firstly designed according to their favorite PAMs of SpCas9 (NGG), SaCas9 (NNGRR), and enAsCas12a (TTTV) systems, respectively. Off-target effects for each gRNA were assessed using the offline version of Cas-Offinder^58^, and gRNAs with multiple binding sites were excluded. Remaining gRNAs were further filtered to pass all the following restrictions: (1) no multiple repeat nucleotide (e.g., TTTT or AAAA); (2) the GC content of the gRNA ranges from 20% to 80%. The selected gRNAs were then assembled into pairs to ensure that the dual-cut fragments fallen within a range of 50–200 bp. For each peak, we randomly selected 35 pgRNA from all gRNA pairs. For the sgRNA-mediated perturbation platforms, all SpCas9-associated gRNAs used in the pgRNA library (dual-SpCas9 and Papi) were included. To ensure that the target regions overlapped with those in the pgRNA platforms, additional gRNAs whose target loci were within the dual-cut fragment regions were also incorporated. In addition, control sgRNAs or pgRNAs targeting the *AAVS1*, *CCR5*, and *ROSA26* safe harbor loci as well as non-targeting ones were included. Three independent pgRNA libraries with 2,480, 2,200 and 747 gRNA pairs were designed for dual-SpCas9, Papi and dual-enAsCas12a, respectively. The sgRNA library with 3,870 gRNAs was designed for both CRISPR KO and CRISPRi platforms.

For the lncRNA library design, we selected lncRNAs that were highly expressed (fold change > 2 & FDR < 0.01) in both lung adenocarcinoma (TCGA-LUAD) and lung squamous cell carcinoma (TCGA-LUSC) tumor samples (n = 533 for LUAD, n = 502 for LUSC), compared with normal tissue samples (n = 59 for LUAD, n = 49 for LUSC). These lncRNAs were mapped to the hg38 reference genome to determine their precise genomic locations. Using such criteria, a total of 220 lncRNA candidates were chosen for perturbation. For the gRNA design, a length of ∼400 bp genomic region flanking the TSS of each lncRNA was selected for targeting. Similar procedures were followed for the gRNA selection as above in the enhancer library design. For pgRNA systems, an additional criterion was applied: the dual-cut fragment of a pgRNA pair was required to span both sides of TSS for the target lncRNA. Control gRNAs targeting the safe harbor loci (*AAVS1*, *CCR5*, and *ROSA26*) and the coding regions of several core essential genes (*EEF1A1*, *EEF2*, *RPL7*, *RPS19*, and *RPS20*) as well as the non-targeting gRNAs were also included. Three independent pgRNA libraries with 3,595, 3,195 and 2,033 gRNA pairs were designed for dual-SpCas9, Papi and dual-enAsCas12a, respectively. The sgRNA library with 4,217 gRNAs was designed for both CRISPR KO and CRISPRi platforms. The DNA oligo libraries were synthesized by Synbio Technologies company (China). The details of the gRNA information in these libraries are provided in Supplementary Table 1 and 3.

### Plasmid library construction

For pgRNA library construction, a two-step strategy was employed.

#### Step I: Generation of the First Plasmid Library Containing pgRNAs

Vectors such as pLentiGuide-Puro (dual-SpCas9), pPapi (Papi), and pRDA-052 (dual-enAsCas12a) were digested with the Esp3I (BsmBI) restriction enzyme (Thermo Fisher Scientific, #FD0454) at 37 °C for 1.5 h. DNA oligo libraries corresponding to the three different pgRNA systems were demultiplexed by PCR, purified using the GeneJET PCR Purification Kit (Thermo Fisher Scientific, K0702), and subsequently digested with BpiI (BbsI) restriction enzyme (Thermo Fisher Scientific, #FD1014) at 37 °C for 1.5 h. The digested vectors and oligo libraries were purified using the GeneJET Gel Extraction Kit (Thermo Fisher Scientific, K0692) and assembled using T4 DNA ligase (Thermo Fisher Scientific, #EL0016) at a 3:1 molar ratio of insert to vector. The assembled plasmid library was then transformed into self-prepared electrocompetent Stable *E. coli* cells (New England Biolabs, #C3040) by electroporation, achieving a transformation efficiency that ensured at least 50X coverage representation of each designed clone. Transformed bacteria were then cultured in liquid LB medium with ampicillin at 30°C for 16 h, followed by plasmid extraction using the Endo-Free Maxi-Prep Plasmid Kit (TIANGEN, #4992194). The library in current form for dual-enAsCas12a system was ready for use without subsequent step.

#### Step II: Generation of the Second Plasmid Library by Inserting the gRNA Scaffold and Promoter

To complete the dual-SpCas9 and Papi library construction, the first plasmid libraries generated in Step I were digested with Esp3I (BsmBI) at 37 °C for 1.5 h. The linearized and dephosphorylated plasmids were separated by 1.0% agarose gel electrophoresis and purified using the GeneJET Gel Extraction Kit. A synthesized H1 promoter and gRNA scaffold sequence was also digested with Esp3I and inserted into the digested dual-SpCas9 first plasmid library using T4 DNA ligase. Similarly, a fragment with two different gRNA scaffold (SpCas9 and SaCas9) sequences was inserted into the first Papi plasmid library following the same procedures. The inserted sequence is listed in Supplementary Table 5. After bacterial transformation and plasmid extraction, the second plasmid libraries for dual-SpCas9 and Papi systems were ready for use. A schematic describing the pgRNA library construction was shown in Extended Data Fig. 3c-e.

For the sgRNA plasmid library construction, lentiCRISPRv2 (CRISPR KO) and plenti-dCas9m4-KRAB-MECP2-puro (CRISPRi) vectors were also digested with Esp3I at 37 °C for 1.5 h, followed by gel purification. Oligo libraries corresponding to CRISPR KO and CRISPRi systems were demultiplexed by PCR and purified. Overlapping sequences with the vector were introduced by PCR, and the purified fragments were assembled into the vector using 2X Gibson Assembly Mix at 50 °C for 1 h. Following assembly, the plasmid libraries were transformed, cultured, and extracted using the same procedures as described above during pgRNA plasmid library construction. A schematic describing the sgRNA library construction was shown in Extended Data Fig. 3a,b.

### Cell culture

HEK293FT, A375, A549, K562, and PC-9 cell lines were obtained from the American Type Culture Collection (ATCC). HEK293FT, A375 and A549 cells were maintained in Dulbecco’s Modified Eagle Medium (DMEM) (VivaCell BIOSCIENCES, #C3114-0500) supplemented with 10% fetal bovine serum (FBS) (ExCell, #FSP500) and 1% penicillin–streptomycin (VivaCell BIOSCIENCES, #C3420-0100). K562 and PC-9 cells were cultured in RPMI-1640 medium (VivaCell BIOSCIENCES, #C3001-0500) with 10% FBS and 1% penicillin–streptomycin. All cell lines were maintained at 37 °C with 5% CO₂ and were routinely tested to confirm the absence of mycoplasma contamination.

### Lentivirus production

HEK293FT cells were seeded in 6-well plates and transfected when reaching 60%– 80% confluency. For each well transfection, the Opti-MEM reduced-serum medium (Gibco, #51200038) was mixed with 5 μL of NeoFect Transfection Reagent, 2 μg of lentiviral vector, 1.5 μg of pCMVR8.74, and 1 μg of pMD2.G, bringing the final volume to 0.2 mL. The mixture was incubated at room temperature for 25 min before adding to the cell culture medium. After 6-12 h post transfection, the medium was replaced with fresh DMEM supplemented with 20% FBS. The lentivirus-containing supernatant was collected at 48 h post transfection, centrifuged at 5,000 rpm for 5 min to remove cell debris, aliquoted, and stored at −80 °C until further use. For large-scale production in 100 mm culture dishes, the same protocol was followed, with all reagent volumes scaled up fivefold.

### CRE or enhancer screens

For each screen, around 5×10^6^ A549 or K562 cells expressing minimally low copies of Cas effectors via low MOI lentiviral infection were transduced with lentiviruses delivering corresponding enhancer-targeting sgRNA or pgRNA libraries at a low MOI (∼0.3). After 2 days, cells were selected with 1 μg/mL puromycin (Solarbio, #B9300) for 3 days followed by additional two-day recovery. At least 2×10^6^ (more than 500X coverage) cells were collected as Day 0 sample, and the remaining cells were divided into either DMSO vehicle or drug treatment (2.5 μM 6TG) groups. Each group contains at least 2×10^6^ cells at Day 0. After three weeks, the cells were collected as the sample of screening endpoint (with at least 2×10^6^ cells for each sample). The genomic DNA was isolated for all the collected samples and the gRNA fragment was PCR-amplified to construct the sequencing libraries. The abundance of the gRNAs was determined by high-throughput sequencing with PE150 mode.

### LncRNA screens

For the five perturbation systems, the general steps of the lncRNA screens were similar to enhancer screens described above except that: (1) PC-9 cells were employed; (2) the sgRNA or pgRNA libraries were lncRNA-targeting libraries; and (3) the cells were normally maintained for three weeks without drug treatment for cell fitness readout during the screening process. The abundance of the gRNAs of the collected samples was determined by high-throughput sequencing with PE150 mode.

### Individual sgRNA or pgRNA insertion into plasmid vector

For sgRNA-dependent perturbation platform (CRISPR KO and CRISPRi), the gRNA sequence synthesis was as follows (5’-3’): Forward (F) sequence: CACCgN_20_(gRNA sequence 5’-3’); Reverse (R) sequence: AAACN_20_(gRNA sequence 3’-5’)C. For dual-SpCas9 platform, the gRNA sequence synthesis was as follows: F-sequence: CACCG(gRNA1 5’-3’)GTTTGGAGACGCGTCTCCCCCG(gRNA2 5’-3’); R-sequence: AAAC(gRNA2 3’-5’)CGGGGGAGACGCGTCTCCAAAC(gRNA1 3’-5’)C. For Papi system, the gRNA sequence synthesis was as follows: F-sequence: CACCG(Sp-gRNA 5’-3’)GTTTGGAGACGCGTCTCCTAAAAC(Sa-gRNA 3’-5’)C; R-sequence: TCCCG(Sa-gRNA 5’-3’)GTTTTAGGAGACGCGTCTCCAAAC(Sp-gRNA 3’-5’)C. For dual-enAsCas12a platform, the gRNA sequence synthesis was as follows: F-sequence: AGAT(gRNA1 5’-3’)TAATTTCTACTCTTGTAGAT(gRNA2 5’-3’)TTTTTTT; R-sequence: ATTCAAAAAA(gRNA2 3’-5’)ATCTACAAGAGTAGAAATTA(gRNA1 3’-5’).

The forward and reversed oligonucleotides were annealed in the following reaction: 10 μM forward sequence oligo, 10 μM reversed oligo, T4 ligation buffer (1X), and 5U of T4 polynucleotide kinase (thermo, #EK0032) with the cycling parameters of 37 °C for 30 min; 95 °C for 5 min and then ramp down to 25 °C at 1 °C/min in a PCR machine. The annealed oligos were then cloned into their corresponding vectors by a T4 ligation reaction with a mix of 100 ng of Esp3I linearized vector, 0.1 μM annealed oligos, T4 ligation buffer (1X), and 5U of T4 DNA ligase (thermo, #EL0016). Incubate the mix at room temperature for 1.5 h, transform the Stable *E. coli* cells and extract the plasmids using the EndoFree Mini Plasmid Kit II (TIANGEN, #DP118-02). The individual gRNA sequences are listed in Supplementary Table 5. The scaffold and promoter sequences were then inserted in the secondary cloning step to obtain the final pgRNA constructs.

### Western blot

For protein extraction, cells were lysed in ice-cold RIPA lysis buffer (Beyotime, # P0013C), supplemented with a Protease Inhibitor Cocktail (CWBIO, #CW2200S), and incubated on ice for 20 min. The protein lysate was then centrifugated at 14,000 rpm for 10 min at 4 °C, and the supernatant was collected. Equal amounts of protein were mixed with 5X loading buffer and denatured by boiling at 95 °C for 5 min. Protein samples were separated by SDS-PAGE, transferred to a PVDF membrane and blotted using the following primary antibodies: FLAG antibody (Invitrogen, #MA1-91878), GAPDH antibody (Santa Cruz, #sc-25778), HA antibody (Proteintech, #51064-2-AP), HPRT1 antibody (Proteintech, #67518-1-lg), PCNA antibody (Beyotime, #AF1363), PMS2 antibody (Beyotime, #AF1339), MLH1 antibody (Beyotime, #AF1327), MHS2 antibody (Beyotime, #AF7509), and MSH6 antibody (Beyotime, #AF0237). The secondary antibodies were Goat Anti-Mouse IgG H&L (Abcam, # ab150116) and Goat Anti-Rabbit IgG H&L (Abcam, # ab150080). The blotting results were visualized by chemiluminescence (Sparkjade, #ED0025) and the signal was recorded using an imaging system (Tanon, #5200).

### Quantitative reverse transcription PCR (RT-qPCR) analysis

To determine relative gene expression at the RNA level, total RNA was extracted from collected cells using the UNlQ-10 Column Trizol Total RNA Isolation Kit (Sangon Biotech, #B511321). For reverse transcription, 1 μg of RNA was incubated with random primers and MultiScribe reverse transcriptase (Thermo Fisher, #4311235) at 25 °C for 10 min, 37 °C for 90 min, and 85 °C for 5 min. Quantitative PCR (qPCR) was then performed using 2X UltraSYBR mixture (Sparkjade, #AH0105) with *RPS28* gene serving as the housekeeping gene for normalization. The qPCR reaction was carried out on a QuantStudio 5 Real-Time PCR System (Thermo Fisher, A28319) with the following program: 10 min at 95 °C for pre-denaturation, followed by 40 cycles of 10 s at 95 °C (denaturation), 30 s at 60 °C (annealing), and 30 s at 72 °C (elongation). Each sample was run in triplicate. Relative gene expression was calculated using the ΔΔCt method. The primers for qPCR are listed in Supplementary Table 5.

### Genomic DNA extraction

All genomic DNA was extracted from collected cells using the lysis buffer (300 mM NaCl, 0.2% SDS, 2 mM EDTA, 10 mM Tris-HCl pH 8.0) supplemented with RNase A (10 mg/mL), followed by mixing and incubation at 65 °C for 1 h. Then the proteinase K (10 mg/mL) were added followed by incubation at 55 °C overnight (or 6 h). The solution was mixed with phenol/chloroform/isoamyl alcohol solution (25:24:1) for 1:1 and centrifuged at 12,000 rpm for 10 min before taking the upper phase. The genomic DNA was precipitated using isopropanol and washed with 75% ethanol before being dissolved in nuclease-free water.

### Dual-cut fragment editing assay

For each target site, the individual pgRNA was lentivirally transduced into Cas effector-expressing cells at a low MOI (∼0.3) for corresponding dual-cut perturbation system. After 7 days (2-day infection + 3-day puromycin selection + 2-day recovery), the genomic DNA was extract from collected cells. Around 300 ng of genomic DNA were used to amplify the dual-cut fragment editing region with Extaq (Takara, #RR001B) using the standard program. The PCR product was separated with native 10% TBE-PAGE gel and stained with 0.5 µg/mL ethidium bromide. The band intensity in the image was analyzed by the ImageJ software. The dual-cut fragment editing efficiency was determined by the editing ratio of Ei to (Ai + Ei), in which Ei stands for edited fragment intensity and Ai means amplified fragment intensity (Fig. 1c). Three biological replicates were performed for each perturbation case. The pgRNA sequences and primers are listed in Supplementary Table 5.

### shRNA knockdown assay

For each target lncRNA, shRNAs were designed using the GPP Web Portal (https://portals.broadinstitute.org/gpp/public/) and cloned into the pLKO.1-puro vector (Addgene, #8453). Individual shRNAs were lentivirally transduced into the cells at a high MOI (>1). Total RNA was extracted 2 days post-infection, and lncRNA expression levels were quantified by RT–qPCR in triplicates. Following the target lncRNA knockdown, around 5,000 cells per well were seeded into 12-well plates, and the cell number was determined using a hemocytometer after 7 days post seeding. Three biological replicates were performed for each perturbation. The shRNA sequences and RT–qPCR primers are listed in Supplementary Table 5.

### Data processing for CRE screens

The sequencing data of CRISPR KO and CRISPRi screens in FASTQ format were processed using MAGeCK^59^ to generate the sgRNA raw read count table. For the dual-SpCas9, Papi, and dual-enAsCas12a screening raw sequences, the pgRNA information was aligned using Bowtie^60^ followed by removing the recombination reads. The recombination read rate for pgRNA samples ranged from 46% - 51%, with the average rate around 49%. Subsequently, a Robust Rank Aggregation (RRA) algorithm was applied to normalize and rank the pgRNA read counts. We then computed the raw log fold change (LFC) value for each sgRNA or pgRNA using the MAGeCK. We then calibrated the LFC values by using the average LFC value of non-targeting gRNAs from the Vehicle (DMSO) group as the background for the *Z*-score calculation across multiple conditions and systems. To better compare the gRNA performance across different perturbation systems, we further employed a sigmoid function^61^ to normalize the *Z*-score between 0 to 1. The calculation formula is as follows:

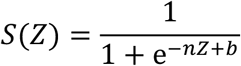

*Z* represents the *Z*-score of LFC for each sgRNA or pgRNA. The n and b are two customized parameters to nominate:

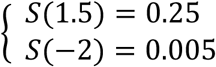

The custom parameters represent a unified processing scheme determined based on the LFC distribution across all the editing systems. The goal was to map invalid LFC values (<−2) in the positive selection screens to a range closer to 0 and the high LFC (strongly positive selection) values to a range closer to 1. The positively selected gRNAs in each system were defined by a sigmoid score > 0.25. The functional (positively selected) CREs in each system were defined with the following criteria: (1) a qualified gRNA with sigmoid score^6TG^ > 0.25 & (score^6TG^ - score^DMSO^) > 0.1; and (2) at least 2 qualified gRNAs exist for the target CRE. The detailed screening results were shown in Supplementary Table 2.

### Data processing for lncRNA screens

The processing of FASTQ reads from the lncRNA screens, including alignment and read count calculation using MAGECK with the Robust Rank Aggregation (RRA) algorithm, was performed following similar procedures as used for the CRE screen analysis. A log10 transformed RRA scores (tRRA) corresponding to different screening directions were obtained using the ReadRRA function implemented in MAGeCK. A higher positive score for a given lncRNA indicates stronger positive selection upon perturbation, while a lower negative score means stronger negative selection.

### Receiver Operating Characteristic (ROC) curve analysis

For the CRE screens, we defined the reference true positive (TP) samples as those high-confidence functional CREs that were shared by at least three out of the five perturbation systems according to the above cutoff. On the other hand, the true negative (TN) samples were defined as those non-functional controls (*AAVS1*, *CCR5* or *ROSA26* sites). For the lncRNA screens, we selected functionally confirmed lncRNAs (supported by published lung cancer studies (Supplementary References) and also identified in our screens by either perturbation system) and non-functional controls (*AAVS1*, *CCR5* or *ROSA26* sites) as a gold standard. Then, the performance of each gRNA in the corresponding perturbation system could be classified as TP, TN, false positive (FP) or false negative (FN). The AUC (Area Under the Curve) was calculated for both types of screens based on the ROC curves.

### Development of DeepDC model

DeepDC is a deep learning-based computational model designed to predict the dual-cut fragment editing efficiency of dual-SpCas9 system. The training data includes 9 published datasets derived from dual-SpCas9 pgRNA-mediated high-throughput screens^16, 17, 19, 20, 22, 62–65^ as well as our CRE screening data generated in this work. To construct DeepDC, we used XGBoost as the underlying architecture, and incorporated gRNA performance score and multiple features (e.g., fragment Hyena score, GC content, length, sgRNA efficiency and epigenetic marks) for the model building. The details were as follows:

#### sgRNA efficiency

To obtain the predicted efficiency score for each sgRNA within the pgRNA pair, multiple scoring tools for sgRNA performance were employed, including DeepSpCas9^63^, DeepCRISPR^66, 67^, CRISPRedict^68^, and Ruleset2^69^. For each pgRNA pair, the harmonic mean of the two single-edit sgRNA scores was calculated as a predictor of the fragment editing efficiency.

#### HyenaDNA score for the edited fragment of dual-cut

HyenaDNA is an autoregressive model based on a decoder architecture^44^. Its core component, the Hyena Operator, replaces the attention mechanism by combining long convolutions with gating. The model was pre-trained on the “Human Regulatory Ensembl” dataset from the Ensembl database, using a learning rate of 6e-4, a decay rate of 0.1, and 128 hidden states over 50 epochs. The best-performing model on the validation set was selected for scoring. The HyenaDNA score for a sequence 𝑆 is calculated as:

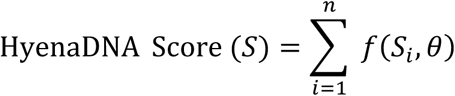

where 𝑓(𝑆*_i_*, 𝜃) represents the score for each nucleotide 𝑆_&_, 𝜃 is the full set of learned parameters of the model (weights of Hyena Operator, gating mechanisms, decoder layers, etc.) obtained from training. These parameters determine how the model assigns scores to each nucleotide and 𝑛 is the sequence length.

#### Calculation of epigenetic marks

DeepDC utilized epigenetic data, including DNase-seq, ATAC-seq, H3K27ac ChIP-seq, and H3K4me3 ChIP-seq, with bigWig datasets from ENCODE (https://www.encodeproject.org/) and GEO database (https://www.ncbi.nlm.nih.gov/geo/). Sequences were aligned to the hg38 human reference genome using Bowtie. We applied TPM (Transcripts Per Million) normalization to address dataset quality imbalances across samples.

#### Sequence collection and processing

The pgRNA sequences of the dual-SpCas9 system along with the target region served as input sequences, which were converted into a four-dimensional binary matrix using one-hot encoding. The edited fragment sequences were processed by extracting full sequences from the reference genome and sequence scores were obtained using HyenaDNA.

#### XGBoost model training and evaluation

An XGBoost regression model was employed to predict continuous outcomes ranging from 0 to 1. The training data (𝑋_train_ 𝑦_train_) and testing data (𝑋_test_, 𝑦_test_) were converted into DMatrix objects for efficient processing. The model was trained using the following hyperparameters: a regression objective function, RMSE as the evaluation metric, a learning rate (𝜂) of 0.05, a maximum tree depth of 20, subsample and column sample rates of 0.8, and training on the CPU. The model was trained for 80 rounds, with training time recorded for efficiency assessment. After training, predictions were made on the test set and evaluated using Root Mean Squared Error (RMSE) and R-squared (𝑅^)^).

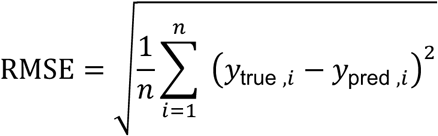

where 𝑛 is the number of samples, 𝑦_true,&_ are the true values, and 𝑦_pred,&_ are the predicted values.

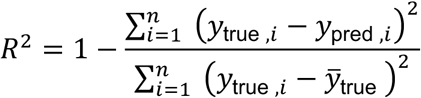

where 𝑦̂_true_ is the mean of the true values.

The formula for the objective function in XGBoost’s regression task is:

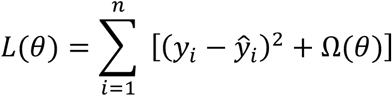

where 𝑦_i_ is the true label, 𝑦^_i_ is the predicted label, and Ω(𝜃) is the regularization term:

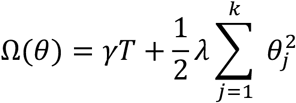

with 𝑇 being the number of leaves in a tree, 𝜃, the parameters for the 𝑗-th feature, and 𝛾, 𝜆 the regularization hyperparameters. After selecting the optimal hyperparameters, we used the full training dataset with selected hyperparameters to train the final model.

### Performance comparison of DeepDC with existing methods

We compared the performance of DeepDC with other published sgRNA efficiency predictor models, including DeepCRISPR^62^, DeepSpCas9^63^, DeepCpf1^66, 67, 70^ and the Random Forest algorithm. For all methods used in the comparison, we provided the same training and test datasets and evaluated them using their recommended default parameters.

### Building the web server

The current version of DeepDC was developed using MySQL 8.0 (http://www.mysql.com) and maintained on a Linux-based Nginx server. We used Python 3.8 and Tornado 6.1 (https://www.tornadoweb.org/) for server-side scripting, including all the interactive functions. The interactive frontend interface was designed and built using vue 2 (https://v2.vuejs.org/) and element UI (https://element.eleme.io/).

### Statistical analysis

To proceed with statistical analysis, more than three samples or repeats were performed. Data points were shown in the graphs. Error bars represent standard deviation, unless otherwise noted. Statistical considerations are reported in each figure legend. *P* < 0.05 was considered statistically significant.

## Data availability

The data supporting the results in this study are available within the paper and its Supplementary Information. The raw sequencing data generated in this study have been deposited and are publicly available in the Genome Sequence Archive in National Genomics Data Center, China National Center for Bioinformation / Beijing Institute of Genomics, Chinese Academy of Sciences, under accession code (GSA-Human: HRA013783) at https://ngdc.cncb.ac.cn/gsa-human.

## Code availability

The custom code used in this study is available at https://github.com/xmuhuanglab/DeepDC. The DeepDC webserver is available at https://deepdc.huanglabxmu.com/.

## Supporting information

Supplementary References

## Acknowledgements

This work was supported by the National Natural Science Foundation of China (32470673), Guangdong Basic and Applied Basic Research Foundation (2023A1515140084), the 111 Project (B16009) and the Construction Project of Liaoning Provincial Key Laboratory, China (2022JH13/10200026) to T.F., the National Natural Science Foundation of China (92474104, 32370586) to J.H., and the Fundamental Research Funds for the Central Universities (20720230068) to J.H.

## Author contributions

T.F. and J.H. conceived the study. T.F., J.H., H.Z., and S.L. designed the research. H.Z. performed most of the experiments with the help of X.W., C.Z., Y.Z., W.Z., Z.C and X.L. S.L. conducted most of the bioinformatics analysis with the help of L.L., R.L., F.C. and N.S. All the authors analyzed the data. T.F., J.H., H.Z. and S.L. wrote the manuscript with the input from all the other authors. T.F. and J.H. supervised the study.

---

**Extended Data Fig. 1.**
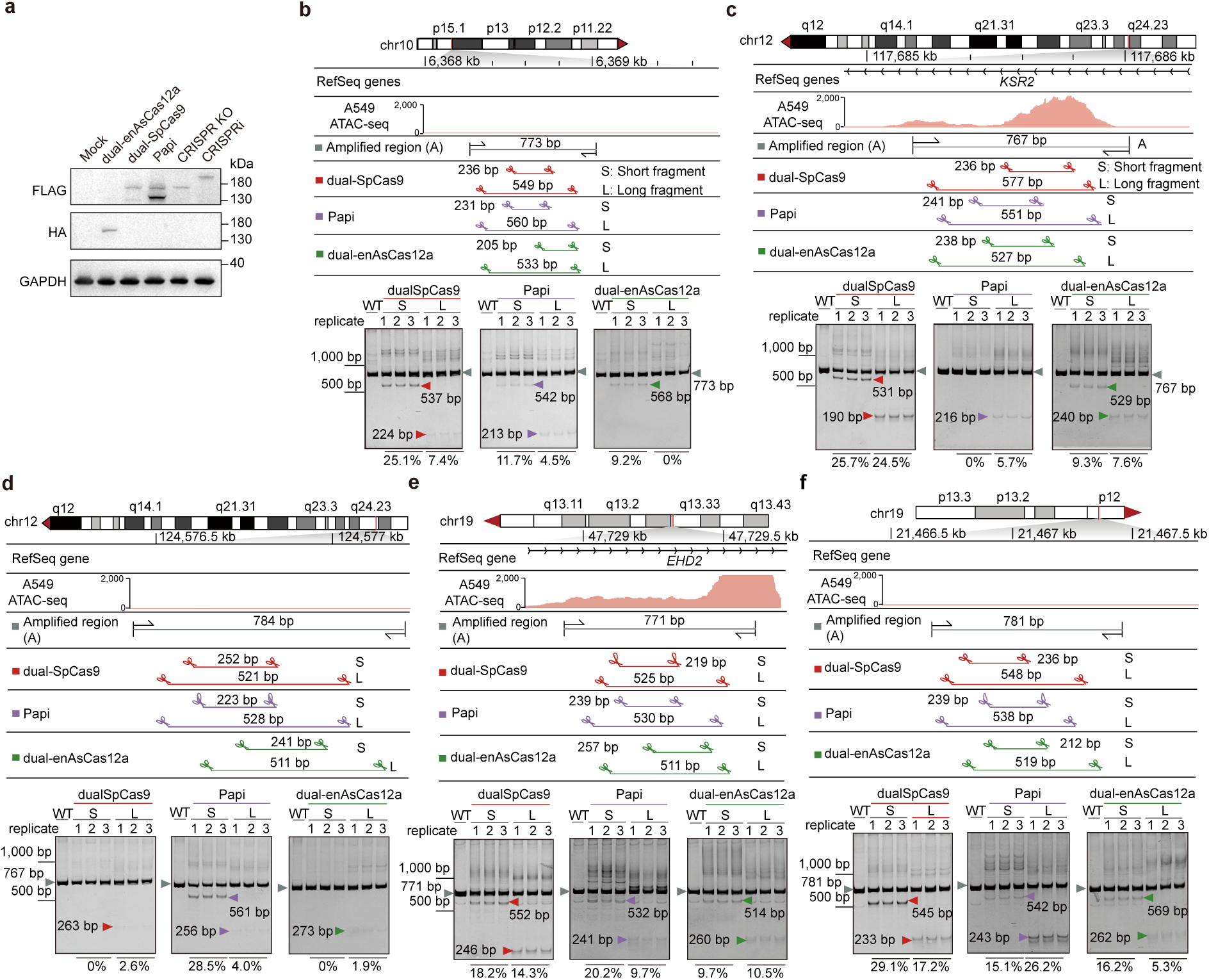
Dual-cut efficiency assays by the three pgRNA-mediated perturbation systems. **a**, Western blot showing the protein expression of lentivirally delivered Cas effectors of the five perturbation systems in A549 cells. The two bands in Papi sample indicate SpCas9 (upper) and SaCas9 (lower), respectively. **b-f**, Dual-cut fragment deletion assay for the indicated pgRNAs to determine the dual-cut editing efficiency in A549 cells at indicated genomic locus. The designed positions of pgRNAs are shown. Edited samples by corresponding perturbation systems are PCR-amplified across the same region and analyzed by native 10% PAGE in TBE (n = 3 biological replicates). The calculated editing efficiency is indicated below the gel image.

**Extended Data Fig. 2.**
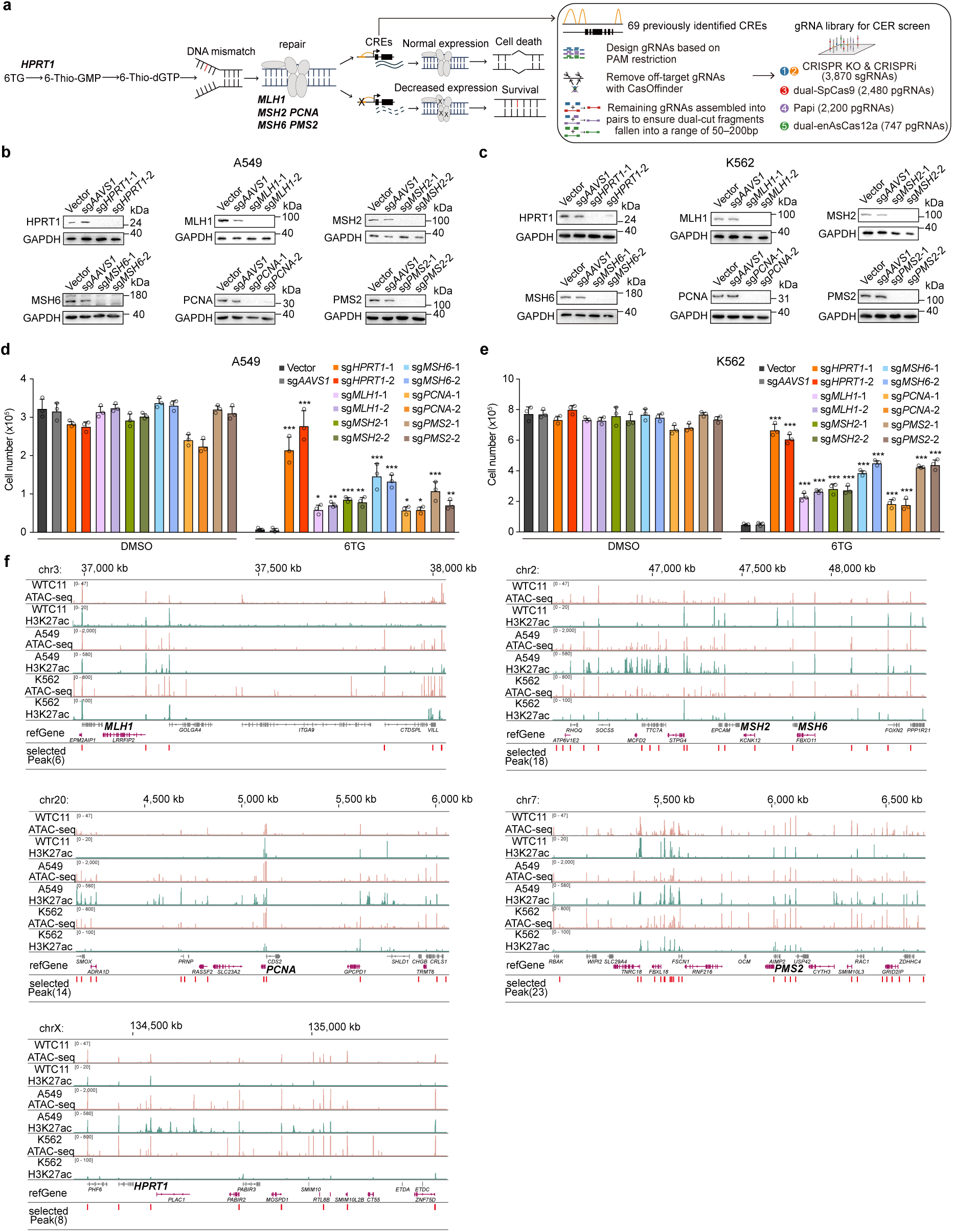
Functional assessment of MMR genes and features of targeted CRE loci in the benchmarking libraries. **a**, Schematic showing the roles of the six genes (*HPRT1, MSH2, MSH6, MLH1, PMS2,* and *PCNA*) in the 6TG-induced mismatch repair (MMR) process and the cell fitness-based CRE screening principle. **b**,**c**, Western blot of A549 (**b**) and K562 (**c**) cells showing the knockout effects of two independent sgRNAs targeting *HPRT1*, *MLH1*, *MSH2*, *MSH6*, *PCNA*, or *PMS2* genes, respectively, at the protein level. Vector and *AAVS1*-targeting sgRNAs served as controls. GAPDH serves as a loading control. **d**,**e**, Cell growth of A549 (**d**) and K562 (**e**) cells upon DMSO or 2.5μM 6TG treatment in control (Vector, sg*AAVS1*) or MMR gene knockout cells (each by two independent sgRNAs) for 7 days. Mean ± SD, n = 3 biological replicates, compared to sg*AAVS1*, one-way ANOVA \**P* < 0.05, \*\**P* < 0.01, \*\*\**P* < 0.001. **f**, The genomic locations and epigenetic features of all selected CREs for perturbation in the benchmarking libraries. The epigenomic data of WTC11 cells were retrieved from GEO database (GSE166834 and GSE166835), and those of A549 and K562 cells were downloaded from ENCODE (https://www.encodeproject.org/).

**Extended Data Fig. 3.**
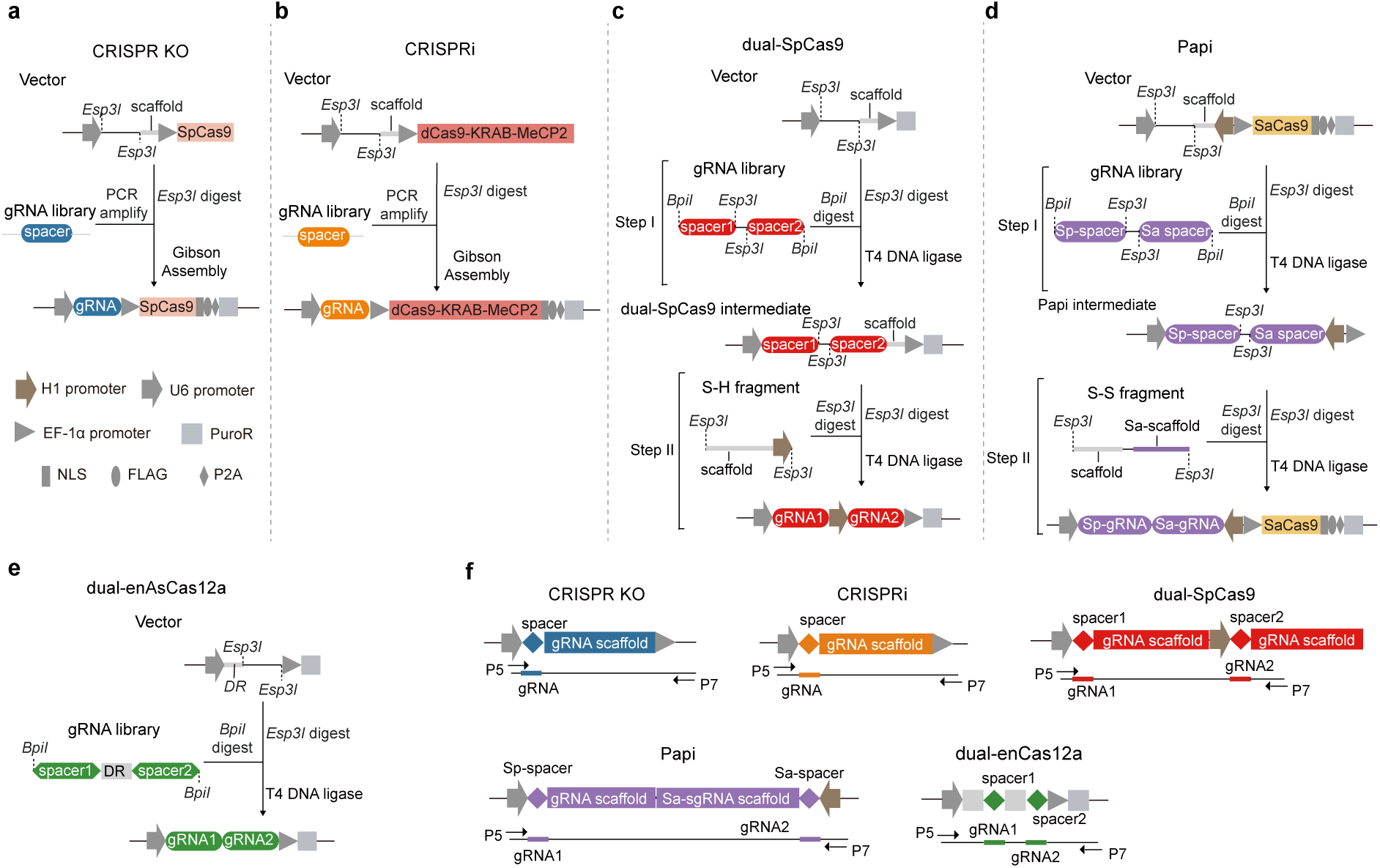
Library construction and sequencing strategies for five perturbation platforms. **a-e**, Schematic representation of library construction process of the five perturbation platforms. **f**, Schematic of next-generation sequencing strategies for the gRNA libraries of the five perturbation platforms.

**Extended Data Fig. 4.**
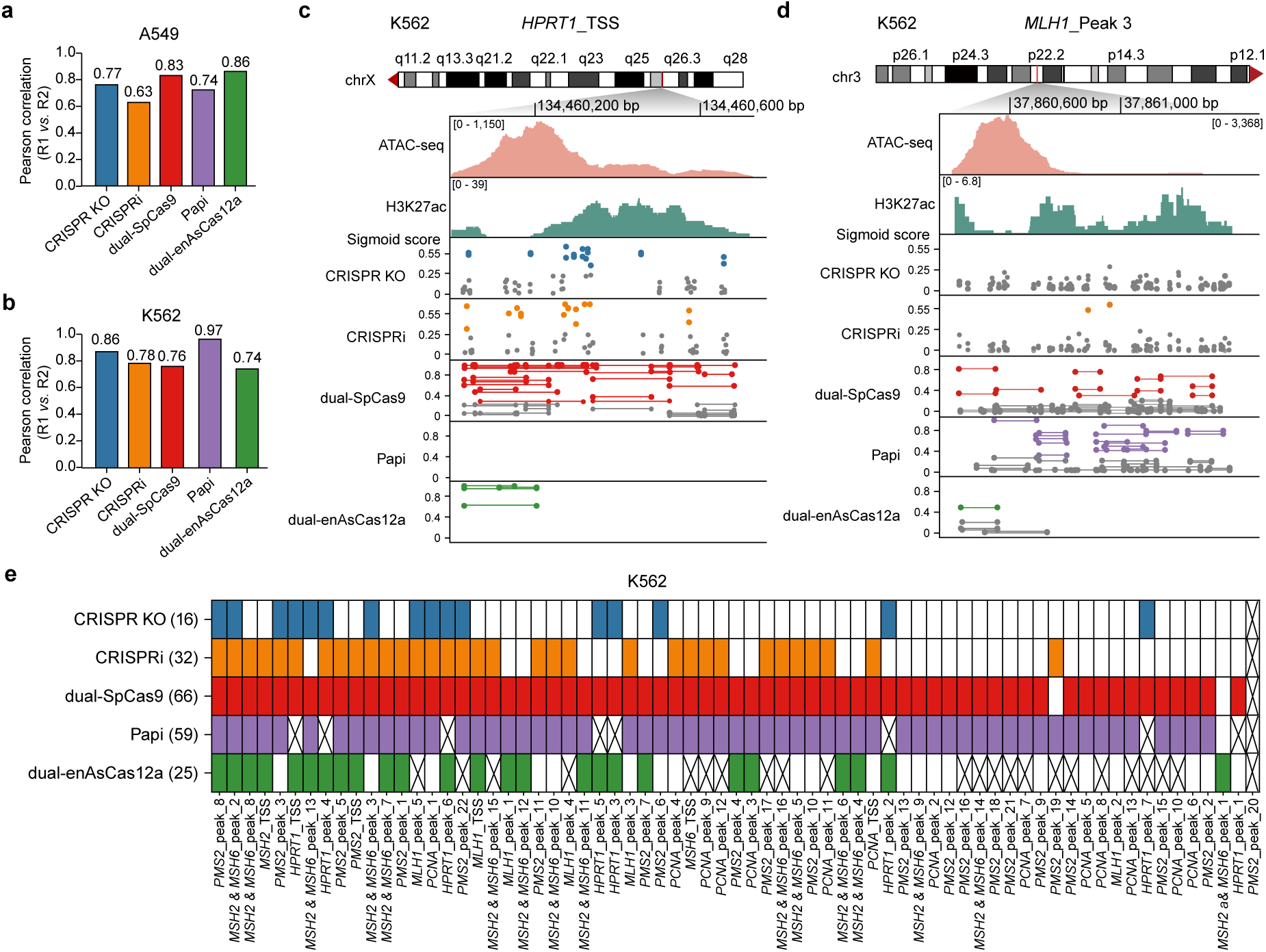
Benchmarking CRE screens in A549 and K562 cells. **a**,**b**, Pearson correlation of the LFC values of TSS-proximal gRNAs in 6TG condition between the two replicates (R1 vs. R2) of CRE screens in A549 (**a**) or K562 (**b**) cells. **c**,**d**, Representative CRE screening results of K562 cells on a TSS region (**c**) or an enhancer (**d**) with gRNA sigmoid score distribution for different perturbation platforms. **e**, Heatmap showing the functional CREs identified in K562 cells by each perturbation platform. The crossed grid indicates that no valid gRNAs were designed for the corresponding CRE.

**Extended Data Fig. 5.**
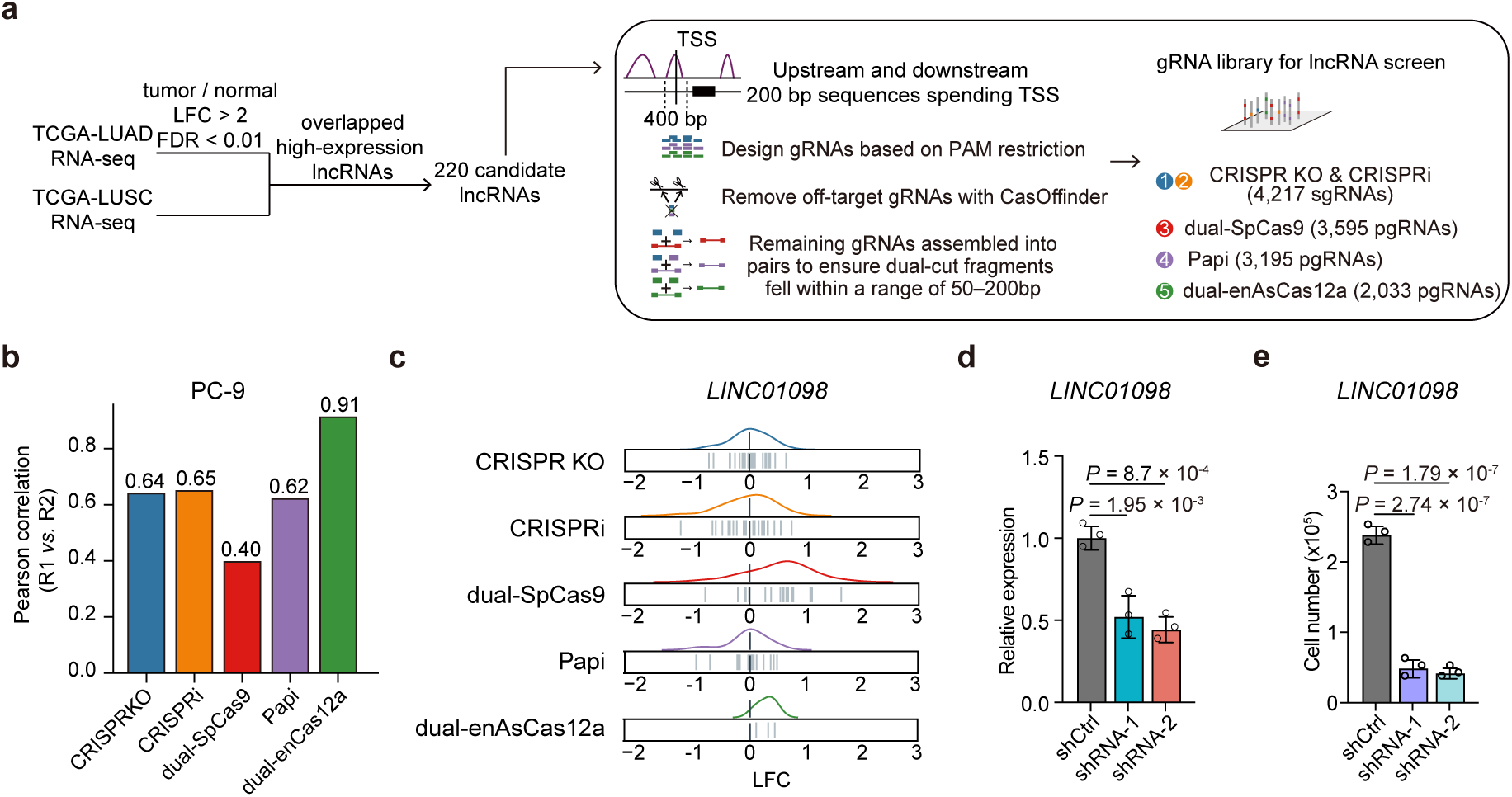
LncRNA screens for benchmarking perturbation tools. **a**, Target lncRNA selection and gRNA library design pipeline. Highly expressed lncRNAs from The Cancer Genome Atlas (TCGA) lung cancer cohorts were selected, and 200 bp upstream and downstream of the TSS region for each lncRNA were chosen for gRNA targeting. Multiple sgRNA and pgRNA libraries were designed. **b**, Pearson correlation of gRNA LFC values between the two replicates (R1 vs. R2) of lncRNA screens in PC-9 cells. **c**, LFC deviation of gRNAs targeting *LINC01098* in the lncRNA screens. **d**, Relative expression of *LINC01098* following shRNA-mediated knockdown in PC-9 cells. Mean ± SD, n = 3, relative to vector control, one-way ANOVA. **e**, Cell growth of PC-9 cells 7 days after shRNA-mediated knockdown of *LINC01098*. Mean ± SD, n = 3 biological replicates, compared to vector control, one-way ANOVA.

**Extended Data Fig. 6.**
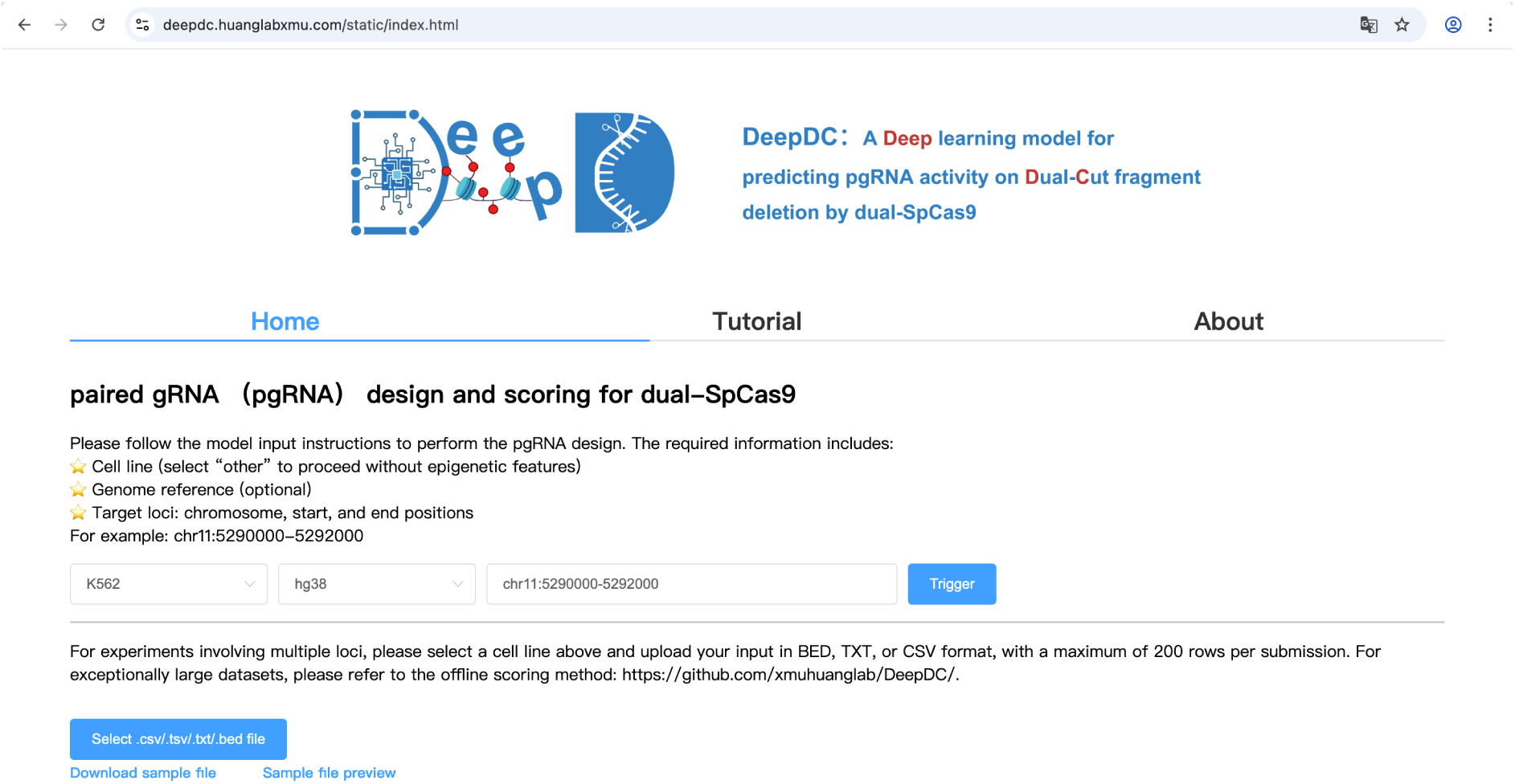
A DeepDC-implemented web server for public access. A snapshot of the frontpage on the DeepDC web server at https://deepdc.huanglabxmu.com/.

